# Exploring the parity paradox: Differential effects on neuroplasticity and neuroinflammation by APOEe4 genotype at middle-age

**DOI:** 10.1101/2023.07.12.548731

**Authors:** Bonnie H Lee, Mel Cevizci, Stephanie E Lieblich, Muna Ibrahim, Yanhua Wen, Rand S Eid, Yvonne Lamers, Paula Duarte-Guterman, Liisa A.M. Galea

## Abstract

Female sex and Apolipoprotein E (APOE) ε4 genotype are top non-modifiable risk factors for Alzheimer’s disease (AD). Although female-unique experiences like parity (pregnancy and motherhood) have positive effects on neuroplasticity at middle age, previous pregnancy may also contribute to AD risk. To explore these seemingly paradoxical long-term effects of parity, we investigated the impact of parity with APOEε4 genotype by examining behavioural and neural biomarkers of brain health in middle-aged female rats. Our findings show that primiparous (parous one time) hAPOEε4 rats display increased use of a non-spatial cognitive strategy and exhibit decreased number and recruitment of new-born neurons in the ventral dentate gyrus of the hippocampus in response to spatial working memory retrieval. Furthermore, primiparity and hAPOEε4 genotype synergistically modulate neuroinflammatory markers in the ventral hippocampus. Collectively, these findings demonstrate that previous parity in hAPOEε4 rats confers an added risk to present with reduced activity and engagement of the hippocampus as well as elevated pro-inflammatory signaling, and underscores the importance of considering female-specific factors and genotype in health research.

Highlights
- hAPOEε4 rats made more errors and used a non-spatial cognitive strategy
- Primiparous hAPOEε4 rats increased use of a non-spatial cognitive strategy
- Parity increased neurogenesis in wildtype rats, but decreased it in hAPOEε4 rats
- Primiparous hAPOEε4 rats had less active new neurons in response to memory retrieval
- Parity and hAPOEε4 affect the neuroimmune milieu in a region-specific manner

---

Alzheimer’s disease (AD) is a progressive neurodegenerative disorder characterized by cognitive decline and pathological changes in the brain (Alzheimer’s Association, 2023). Three well-established non-modifiable risk factors are advanced age, Apolipoprotein E (APOE) ε4 genotype, and female sex (Riedel et al., 2016). Females show greater lifetime risk, more severe neuropathology, and faster cognitive decline associated with AD than males (Barnes et al., 2005; Ferretti et al., 2018; Irvine et al., 2012; Laws et al., 2016; Sohn et al., 2018). Individuals with 1 or 2 APOEε4 alleles have a 3 or 15 fold increased risk of AD, respectively, (Farrer et al., 1997; Neu et al., 2017), and APOEε4 carriers have more severe AD neuropathology, decreased hippocampal volume, and greater memory impairment compared to non-carriers (Manning et al., 2014; Tai et al., 2013; Wolk et al., 2010). Remarkably, APOEε4 genotype confers greater burden of disease particularly for females, as female APOEε4 carriers present with higher lifetime risk and earlier onset of AD, elevated levels of tau pathology, and greater cognitive decline compared to male carriers (Altmann et al., 2014; Duarte-Guterman et al., 2021; Farrer et al., 1997; Hohman et al., 2018; Irvine et al., 2012; Neu et al., 2017). As such, expanding our understanding about the relationship between sex and APOEε4 genotype will be fruitful in advancing the field of AD.

To better understand why females are disproportionately affected by AD, with and without APOEε4, it is essential to understand how female-unique experiences may contribute to AD risk. Previous parity (pregnancy and parenting) influences the trajectory of cognitive and brain aging in humans and rodents (Barha et al., 2015; de Lange et al., 2020; de Lange et al., 2019; Duarte-Guterman et al., 2023; Eid et al., 2019). Although the literature is mixed, studies suggest that parity is associated with an earlier onset of AD (Colucci et al., 2006), and multiparous females (five or more reproductive experiences) have a 1.7-fold increased risk of AD compared to those with fewer reproductive experiences (Jang et al., 2018). Females with two or more reproductive experiences also show greater levels of AD-related neuropathology compared to those with fewer or no children as well as compared to males with children (Beeri et al., 2009). However, the association between parity and dementia risk is not uniform across geographical regions, such that multiparous females showed higher risk in Europe and Latin America, yet nulliparous (never parous) females showed higher risk in Asia (Bae et al., 2020). These discrepancies may be attributed to differences in experiences and susceptibilities related to the ethno-racial, societal, and/or cultural influences across geographical regions. Nonetheless, it is apparent that parity impacts AD risk and pathology. Despite this, the effects of parity on AD-related biomarkers, and further, how APOEε4 genotype may contribute to those relationships, remain unknown, and the current study serves to address these knowledge gaps.

The hippocampus and frontal cortex are among the first areas to be affected in AD (Mueller et al., 2010; Thal et al., 2000). The hippocampus is particularly compelling to investigate as the dentate gyrus of the hippocampus continues to produce new neurons throughout life, and alterations in neurogenesis have been linked to AD-related neuropathology and cognitive impairment in both rodent models and humans (Hollands et al., 2017; Jin, Galvan, et al., 2004; Jin, Peel, et al., 2004; Moreno-Jiménez et al., 2019). In AD mouse models, neurogenesis is reduced with hAPOEε4 genotype, although this has been suggested to be dependent on age and disease progression (Koutseff et al., 2014; Tensaouti et al., 2018). Moreover, patients with amnestic mild cognitive impairment have fewer proliferating neuroblasts than cognitively normal individuals (Tobin et al., 2019), and the number of new-born neurons in the dentate gyrus progressively declines with AD severity (Ekonomou et al., 2015; Moreno-Jiménez et al., 2019). Although some papers have reported no detectable levels of neurogenesis in the human adult hippocampus (Franjic et al., 2022; Sorrells et al., 2018), many more studies using multiple methods show robust evidence of neurogenesis in the adult human hippocampus (Boldrini et al., 2018; Eriksson et al., 1998; Ernst et al., 2014; Gage, 2019; Kempermann et al., 2018; Knoth et al., 2010; Moreno-Jiménez et al., 2019; Sanai et al., 2011; Spalding et al., 2013; Tobin et al., 2019; Zhou et al., 2022).

Interestingly, previous parity has lasting impacts on hippocampal plasticity (Puri et al., 2023). At middle age, primiparous and multiparous rats show greater levels of neurogenesis compared to nulliparous rats (Barha et al., 2015; Eid et al., 2019). In humans, previous parity was associated with younger brain age relative to chronological age, particularly in the limbic and striatal regions, including the hippocampus (de Lange et al., 2020b; de Lange et al., 2019). Collectively, the human and animal research demonstrate that parity has positive long-term effects on neuroplasticity. However, as highlighted earlier, there is also evidence indicating that parity history may confer greater AD risk and pathology (Beeri et al., 2009; Colucci et al., 2006; Jang et al., 2018). One explanation to reconcile these seemingly opposing findings is the Healthy Cell Bias, such that parity may influence AD endophenotypes depending on presence of disease or disease risk. Indeed, Cui et al. (2014) found that wildtype parous mice were quicker than nulliparous mice to solve a spatial working memory task, but in an AD transgenic mouse model (APP23), parous mice took longer than nulliparous mice to solve the task. To date, there is limited research on how parity may differentially influence cognition and hippocampal plasticity depending on APOEε4 genotype and thus, this is the central question investigated in the present study.

Increased neuroinflammation is evident in AD, and prolonged activation of the immune response through activation of microglia and secretion of cytokines can exacerbate the neurodegenerative cascade in AD (Kinney et al., 2018). Neuroinflammation also contributes to neuronal loss and alters neurogenesis (Belarbi et al., 2012; Ekdahl et al., 2003; Leng et al., 2023; M. D. Wu et al., 2012; Zunszain et al., 2012). Further, previous parity is associated with long-term changes in inflammation (Duarte-Guterman et al., 2023; Eid et al., 2019; Haim et al., 2017; Posillico & Schwarz, 2016). For example, middle-aged primiparous rats show lower blood levels of cytokines, such as IFNγ, IL-10, and IL-4, compared to nulliparous rats (Eid et al., 2019). Inflammation reduces neurogenesis in the hippocampus via the kynurenine pathway, as inhibiting activation of the pathway via the KMO receptor ameliorates the detrimental effects of IL-1β on neurogenesis (Kanai et al., 2009; Zunszain et al., 2012). Interestingly, dysregulation of the kynurenine pathway is implicated in both aging and AD, and accumulation of downstream metabolites of the kynurenine pathway amplifies excitotoxicity and degeneration in the brain (Sorgdrager et al., 2019; Zwilling et al., 2011). Parity transiently increases kynurenine pathway metabolites (Duarte-Guterman et al., 2023), but to our knowledge, no research has examined how this may be influenced by APOEε4 genotype. As such, the kynurenine pathway is another target investigated in this study.

Our objective in this study was to investigate the long-term influences of previous parity on biomarkers of brain health and whether these are differentially affected based on APOEε4 genotype. We measured the effects of nulliparity and primiparity on a spatial working memory task that relies on the integrity of the hippocampus and frontal cortex, neurogenesis, inflammation (hippocampus and frontal cortex), and the kynurenine pathway in wildtype and humanized (h) APOEε4 knock-in middle-aged female rats. We expected that primiparous wildtype rats would show evidence of positive biomarkers for brain health (enhanced spatial working memory, greater levels of neurogenesis, and alterations to the neuroinflammatory milieu and kynurenine pathway) compared to nulliparous rats, but that this relationship would be reversed with hAPOEε4 genotype.

## 2. Materials and Methods

### 2.1. Animals

Fifty-one age-matched wildtype and humanized (h) APOEε4 knock-in Sprague Dawley female rats were used. All rats were maintained on a 12-hour light/dark cycle (lights on at 07:00 h) in standard laboratory conditions (21 ± 1°C; 50 ± 10% humidity) and given *ad libitum* access to food (Purina Rat Chow) and water. Rats were randomly assigned to parity groups (nulliparous or primiparous).

Nulliparous rats had no sexual or reproductive experience and no exposure to pups. Primiparous rats had one reproductive experience at 4 months of age. Rats were pair-housed except throughout pregnancy and postpartum in primiparous rats – during which nulliparous rats were also single-housed, but in a separate colony room. All experiments were conducted in accordance with ethical guidelines set by the Canada Council for Animal Care, and all procedures were approved by the University of British Columbia Animal Care Committee. All efforts were made to reduce the number and suffering of animals.

### 2.2. Breeding

Primiparous rats were bred starting at 4 months (see Figure 1 for a timeline). During breeding, 2 females and 1 male were paired at approximately 17:00 h and left overnight. Female rats were vaginally lavaged every morning to assess for the presence of sperm. Upon identification of sperm, female rats were considered pregnant, weighed, and single-housed. One day after birth, litters were culled to 5 males and 5 females. Rats were left undisturbed except for cage changing and weighing, which occurred once a week throughout gestation. Maternal behaviours were observed twice a day (once at 09:00 h and once at 17:00) from postpartum day 2 through 8. Each observation lasted 10 minutes and the amount of time spent nursing (including arched-back, blanket, and passive nursing), and off-nest (including sleeping) were recorded. Pups were weaned on postpartum day 23, after which all rats were pair-housed again and left undisturbed until middle age (12-13 months of age).

**Figure 1.**
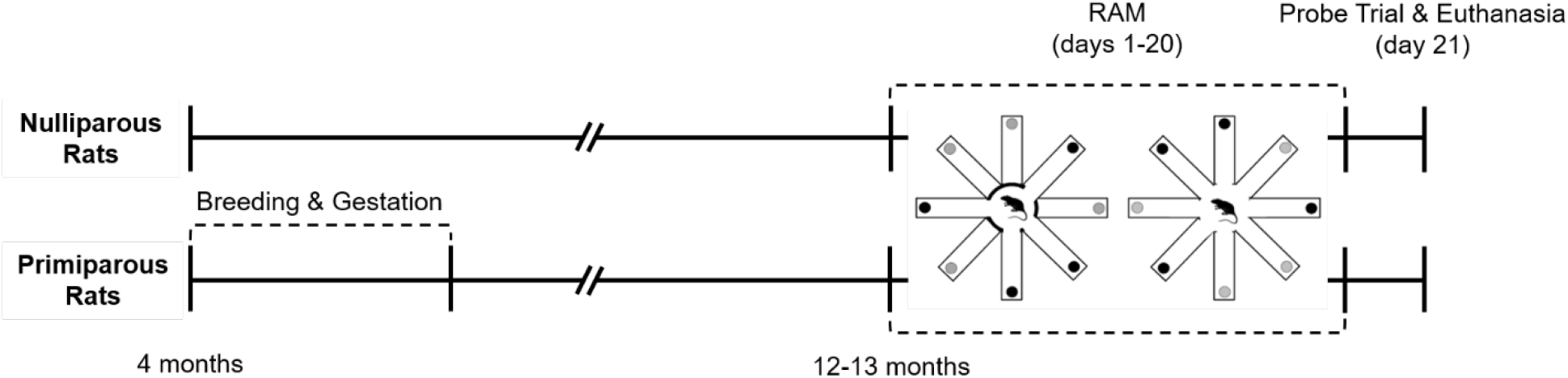
Experimental outline. Wildtype and hAPOEε4 rats were either nulliparous or primiparous. Primiparous rats were bred starting at 4 months. At middle age (12-13 months), all rats were subject to training and testing in the delayed win-shift version of the radial arm maze, then euthanasia.

### 2.3. Apparatus

Rats were trained and tested on a radial arm maze with 8 arms (each 53 cm long x 10 cm wide) extending from a platform in the center (36 cm in diameter), elevated 80 cm above the floor in a dimly lit room. Removable metal barriers were used to block the arms from the center platform. Large extra-maze cues were placed on all four walls of the room and kept constant throughout the experiment.

### 2.4. Procedure

At middle-age (approximately 12-13 months old), rats were single-housed, handled (2-3 minutes per day), and food restricted to 90% of their free-feeding body weight for one week. Then, rats underwent 2 habituation trials followed by 3 shaping trials (1 trial per day across 5 days) (Yagi et al., 2016). Following the last shaping trial, rats were tested in the delayed win-shift version of the radial arm maze. The delayed win-shift task involves daily trials composed of 2 phases (the training phase and the testing phase), for 20 days. During the training phase, 4 of 8 arms of the maze were randomly selected each day and blocked, and the remaining 4 arms were baited with ¼ of a Froot Loop©. Rats were placed in the center of the maze and allowed to explore until all food rewards were eaten or 5 minutes has elapsed. Then, rats were returned to their home cages for a 5-minute delay before the testing phase began. During the testing phase, all arms remained open and the 4 arms that were previously blocked during the training phase were baited with ¼ of a Froot Loop©. Rats were returned to the center of the maze and allowed to explore until all food rewards were eaten or 5 minutes has elapsed. After 20 days of the task, rats were subject to a probe trial on day 21, where everything was kept the same except the delay in between phases was extended to 30 minutes. The maze was cleaned with 70% ethanol between training and testing phases and between rats. The order in which rats were tested was randomized daily. All testing occurred between 11:00 h and 16:00 h.

During both the training and testing phases, latency to enter the first arm and to complete the phase, pattern of arm entries, and whether food rewards were eaten, were recorded. An arm entry was defined as a rat crossing more than halfway down the length of an arm. Errors were scored as across-phase errors, defined as the number of entries into arms baited from training during the testing phase, and within-phase errors, defined as the number of re-entries into any arms during the testing phase. We also investigated strategy use during task acquisition by scoring the number of consecutive entries into adjacent arms and normalizing to the total number of entries.

### 2.5. Tissue and blood collection

Ninety minutes after the probe trial on day 21 of training, rats received a lethal overdose of sodium pentobarbital. Blood was collected via cardiac puncture into cold EDTA-coated tubes and centrifuged 4 hours later for 10 minutes at 4°C, then stored at −80°C. Adrenals were extracted and weighed. Brains were extracted and cut longitudinally into two halves. The right hemispheres were flash frozen on dry ice and stored in −80°C and the left hemispheres were fixed for 24 hours in 4% paraformaldehyde (4°C), then transferred to a 30% sucrose solution for cryoprotection until the brains sank.

### 2.6. Brain tissue processing

The right hemisphere of each brain was sliced into 300μm coronal sections at −10°C using a cryostat (CM3050 S; Leica, Nuβloch, Germany). Punches of 0.75mm and 1.25mm were used to extract tissue from the frontal cortex, dorsal hippocampus, ventral hippocampus, and cortex. The tissue was then homogenized using an Omni Bead Ruptor (Omni International, Kennesaw, GA) with 120μl of cold Tris lysis buffer. Homogenates were centrifuged at 1000xg for 5 minutes at 4°C and supernatants were stored at −80°C for later analysis (Sections 2.6.1 and 2.6.2).

The left hemisphere of each brain was sliced into 35μm coronal sections using a freezing microtome (2M2000R; Leica, Richmond Hill, ON). Sections were collected in series of 5 throughout the frontal cortex and in series of 10 throughout the entire rostral-caudal extent of the hippocampus, and stored in a cryoprotective medium (consisting of 0.1 M PBS, 30% ethylene glycol, and 20% glycerol) at - 20°C. All immunohistochemical procedures (Sections 2.6.3-6) were conducted on free-floating brain sections and on a rotator at room temperature unless otherwise noted. After staining, sections were mounted onto glass slides, allowed to dry, then dehydrated in increasing graded ethanol, defatted with xylene, and cover-slipped with Permount (Fisher Scientific), unless otherwise noted.

#### 2.6.1. Amyloid-beta peptide and phosphorylated tau

To validate the hAPOEε4 rat model used in this research, Aβ42/Aβ40 ratio in the cortex was measured as a proxy for amyloid accumulation in the brain. Compared to Aβ42 or Aβ40 alone, Aβ42/Aβ40 ratio has been demonstrated to be a better reflection of brain amyloid production and its downstream effects, as well as more reliable in accurately diagnosing AD (Dumurgier et al., 2015; Kwak et al., 2020; Lehmann et al., 2018). Amyloid-beta (Aβ) peptide concentrations in the cortex homogenates were measured using a 3-plex electrochemiluminescence immunoassay kit (V-PLEX Aβ Peptide Panel 1) from Meso Scale Discovery (Rockville, MD, USA) according to the manufacturer’s instructions. Samples were run in duplicates to quantify Aβ38, Aβ40, and Aβ42. Tau concentrations in the cortex homogenates were measured using a 2-plex electrochemiluminescence immunoassay kit (V-PLEX Phospho(Thr231)/Total Tau Panel) from Meso Scale Discovery (Rockville, MD, USA) according to the manufacturer’s instructions. Samples were run in duplicates to quantify phosphorylated tau (p-tau) and total tau (tau). Plates were read using a Sector Imager 2400 (Meso Scale Discovery), and data analyses were conducted using the Discovery Workbench 4.0 software (Meso Scale Discovery).

Total protein levels in cortex homogenates were quantified using a Pierce Microplate BCA Protein Assay Kit (Thermo Scientific Pierce Protein Biology, Thermo Fisher Scientific, Waltham, MA, USA) according to the manufacturer’s instructions. Samples were run in triplicates. Aβ peptide and tau levels were normalized to total protein levels and reported as pg/mg of protein.

#### 2.6.2. Cytokines

Cytokine concentrations in the frontal cortex, dorsal hippocampus, and ventral hippocampus homogenates were measured using a multiplex electrochemiluminescence immunoassay kit (V-PLEX Proinflammatory Panel 2, Rat) from Meso Scale Discovery (Rockville, MD, USA) according to the manufacturer’s instructions. Samples were run in duplicates to quantify interferon-gamma (IFN-γ), interleukin-1-beta (IL-1β), interleukin-4 (IL-4), interleukin-5 (IL-5), interleukin-6 (IL-6), interleukin-10 (IL-10), interleukin-13 (IL-13), tumour necrosis factor-alpha (TNF-α), and chemokine (CXC-motif) ligand 1 (CXCL1). Plates were read using a Sector Imager 2400 (Meso Scale Discovery), and data analyses were conducted using the Discovery Workbench 4.0 software (Meso Scale Discovery). This panel consists of a broad range of cytokines, including some traditionally considered proinflammatory (IL-1β, IFN-γ, TNF-α), anti-inflammatory (IL-4, IL-10), and pleiotropic (IL-6), in addition to the chemokine CXCL-1, which is important for neutrophil recruitment. Together, these markers provide a comprehensive assessment of the inflammatory milieu in the brain.

Total protein levels in homogenates from the frontal cortex, dorsal hippocampus, and ventral hippocampus were quantified using a Pierce Microplate BCA Protein Assay Kit (Thermo Scientific Pierce Protein Biology, Thermo Fisher Scientific, Waltham, MA, USA) according to the manufacturer’s instructions. Samples were run in triplicates. Cytokine levels were normalized to total protein levels and reported as pg/mg of protein. T-helper (Th)1 to Th2 ratios were calculated by combining the z-scores of Th1-related cytokines (IL-1β, IFN-γ, TNF-α) and Th2-related cytokines (IL-4, IL-5, IL-10, IL-13) (Berger, 2000; Town et al., 2002).

#### 2.6.3. SRY-box transcription factor 2

One series of hippocampal sections was stained for SRY-box transcription factor 2 (Sox2), which is critical for maintaining pluripotency of radial glia-like cells, and an established marker of neural stem cells and progenitor cells (Amador-Arjona et al., 2015; Berg et al., 2018; Steiner et al., 2006). Tissue was thoroughly rinsed (3 x 10 minutes) in 0.1M tris-buffered saline (TBS; pH 7.4) before staining and between each of the following procedures. Tissue was first treated with 3% hydrogen peroxide (H_2_O_2_ in dH_2_O) for 30 minutes, then blocked with TBS+ solution containing 3% normal horse serum and 0.3% Triton-X in 0.1M TBS for 30 minutes. Tissue was incubated in a primary antibody solution containing 1:1000 mouse anti-Sox2 (Santa Cruz Biotechnology, Santa Cruz, CA, USA) in TBS+ for 48 hours at 4°C. Next, tissue was incubated in a secondary solution containing 1:2 ImmPRESS® (peroxidase) polymer horse anti-mouse IgG (rat absorbed) (Vector Laboratories) in TBS for 30 minutes. This ImmPRESS polymerized reporter enzyme staining system was used to enhance detection of mouse primary antibodies on rat tissues that may contain endogenous rat immunoglobulins. Immunoreactants were visualized with a Nickel-enhanced DAB reaction (Vector Laboratories).

#### 2.6.4. Doublecortin

One series of hippocampal sections was stained for doublecortin (DCX), a microtubule-associated protein expressed in immature neurons. Tissue was thoroughly rinsed (3 x 10 minutes) in 0.1M phosphate-buffered saline (PBS; pH 7.4) before staining and between each of the following procedures. Tissue was first treated with 0.6% hydrogen peroxide (H_2_O_2_ in dH_2_O) for 30 minutes, then incubated in a primary antibody solution containing 1:1000 goat anti-doublecortin (Santa Cruz Biotechnology, Santa Cruz, CA, USA) in 3% normal rabbit serum and 0.4% Triton-X in 0.1M PBS for 24 hours at 4°C. Next, tissue was incubated in a secondary antibody solution containing 1:500 biotinylated rabbit anti-goat (Vector Laboratories, Burlington, ON, Canada) in 0.1M PBS for 24 hours at 4°C. Lastly, tissue was transferred to an avidin-biotin complex (Elite kit; 1:1000, Vector Laboratories) for 4 hours. Immunoreactants were visualized with a Nickel-enhanced DAB reaction (Vector Laboratories).

#### 2.6.5. Doublecortin and zinc finger-containing transcription factor 268

One series of hippocampal sections was double stained for DCX and zinc finger-containing transcription factor 268 (zif268), an immediate early gene (IEG) required for long-term potentiation and memory consolidation (Bozon et al., 2003). Tissue was thoroughly rinsed (3 x 10 minutes) in 0.1M PBS (pH 7.4) before staining and between each of the following procedures. Tissue was first blocked with blocking solution containing 3% normal donkey serum and 0.3% Triton-X in 0.1M PBS for 30 minutes.

Tissue was incubated in primary antibody solution containing 1:500 goat anti-DCX (Santa Cruz Biotechnology, Santa Cruz, CA, USA) and 1:1000 rabbit anti-zif268 (Egr-1 SC-189; Santa Cruz Biotechnology, Santa Cruz, CA, USA) in a dilution solution containing 1% normal donkey serum and 0.3% Triton-X in 0.1M PBS for 24 hours at 4°C. Next tissue was blocked with blocking solution for 30 minutes, then incubated in secondary antibody solution containing 1:500 donkey anti-goat Alexa Fluor 488 (Invitrogen) and 1:500 donkey anti-rabbit Alexa Fluor 594 (Vector Laboratories) in a dilution solution containing 1% normal donkey serum and 0.3% Triton-X in 0.1M PBS. Lastly, sections were mounted onto slides and cover-slipped with PVA DABCO.

#### 2.6.6. Ionized calcium-binding adaptor molecule 1

One series of hippocampal sections was stained for ionized calcium-binding adaptor molecule 1 (Iba1), a wildly used marker of microglia (Korzhevskii & Kirik, 2016). Tissue was thoroughly rinsed (3 x 10 minutes) in 0.1M TBS (pH 7.4) before staining and between each of the following procedures. Tissue was first treated with 3% hydrogen peroxide (H_2_O_2_ in dH_2_O) for 30 minutes, then blocked with TBS+ solution containing 3% normal donkey serum and 0.3% Triton-X in 0.1M TBS for 30 minutes. Tissue was incubated in a primary antibody solution containing 1:1000 rabbit anti-Iba1 (Wako, Osaka, Japan) in TBS+ for 48 hours at 4°C. Next, tissue was incubated in a secondary antibody solution containing 1:250 donkey anti-rabbit Alexa Fluor 594 (Vector Laboratories) in TBS+ for 4 hours. Lastly, tissue was incubated in 1:1000 DAPI in TBS for 2.5 minutes, then mounted onto slides and cover-slipped with PVA DABCO.

### 2.7. Microscopy and cell quantification

An investigator blinded to experimental conditions quantified Sox2-, DCX-, zif258-, and Iba1-immunoreactive (IR) cells. Sox2 sections were imaged with Zeiss Axio Scan.Z1 (Carl Zeiss Microscopy, Thornwood, NY, USA) at 20x magnification using brightfield imaging. Sox2-IR cells were counted automatically from images of 4 sections each from the dorsal and ventral dentate gyrus using a MATLAB (MathWorks) code developed by JEJS and modified by NT as described in Yagi et al. (2020). DCX sections were counted under a 100x objective on an Olympus CX22LED brightfield microscope. DCX-IR cells in the granule cell layer were exhaustively counted from 2 sections each from the dorsal and ventral dentate gyrus. DCX/zif268 sections were imaged with Zeiss Axio Scan 7 (Carl Zeiss Microscopy, Thornwood, NY, USA) at 20x magnification using fluorescent imaging. The percentage of DCX-IR cells co-expressing zif268 were calculated by identifying, exhaustively, whether DCX-IR cells were co-labelled with zif268 from images of 2 sections each from the dorsal and ventral dentate gyrus. Iba1 sections were imaged with Zeiss Axio Scan.Z1 (Carl Zeiss Microscopy, Thornwood, NY, USA) at 40x magnification using fluorescent imaging. Iba1-IR cells were counted from images of 2 sections each from the dorsal and ventral dentate gyrus using ImageJ.

Any codes used can be made available by contacting the corresponding author. Briefly, images are converted to 8-bit, binarized, and thresholded. Size restrictions remove artifacts that are too large or too small to be a cell. The number of times the background is removed from the image is adjusted for each stain to compensate for differences in staining intensity.

### 2.8. Kynurenine pathway plasma metabolite quantification

Plasma samples were thawed and used to quantify tryptophan (TRP), kynurenine (KYN), kynurenic acid (KYNA), xanthurenic acid (XA), anthranilic acid (AA), 3-hydroxykynurenine (3-HK), 3-hydroxyanthranilic acid (3-HAA), and riboflavin (vitamin B2) and its cofactor, flavin adenine dinucleotide (FAD) via isotope dilution liquid chromatography coupled with tandem mass spectrometry, based on a method modified from Midttun et al. (2005). To examine activation of the kynurenine pathway, ratio of kynurenine to tryptophan concentration was calculated.

### 2.9. Estrous cycle classification

Starting at middle-age (12-13 months old), rats were vaginally lavaged daily until euthanasia. Lavage samples were transferred onto microscope slides, stained with Cresyl Violet, and left to dry. Lavage samples were qualitatively categorized as diestrus (consisting primarily of leukocyte-dense cell samples), proestrus (consisting of primarily nucleated epithelial cells), or estrus (consisting primarily of cornified cells), for evidence of irregular cycling (consecutive lavage cycles varying in length and/or order).

### 2.10. Statistical Analyses

Analyses were conducted using Statistica (Statsoft Tulsa, OK). Analysis of variance (ANOVA) were used on most dependent variables of interest with genotype (wildtype, hAPOEε4) and parity (nulliparous, primiparous) as between-subjects variables. For some analyses, repeated measures ANOVA were used with training blocks (i.e. days grouped into 5-day blocks) or region (dorsal and ventral dentate gyrus, frontal cortex) as within-subjects variables. Post-hoc tests used Newman-Keuls and any a priori comparisons were subjected to Bonferroni correction. Any statistical outliers (+/-2 standard deviations group the group mean) were removed. Pearson product-moment correlations were performed between variables of interest. A Chi-square test was used to compare the frequency of rats that had regular estrous cycles. Significance level was set at α=0.05. Effect sizes were calculated as partial η^2^ or Cohen’s d where appropriate.

## 3. Results

### 3.1. hAPOEε4 rats had higher Aβ42/Aβ40 than wildtype rats. Primiparous rats were more likely to display regular estrous cycling than nulliparous rats and there were no group differences in maternal care behaviour, body weight, or adrenal weight

Ratio of Aβ42 to Aβ40 in the cortex was analyzed to confirm genotype differences in amyloid pathology. Indeed, hAPOEε4 rats had a higher Aβ42/Aβ40 than wildtype rats (main effect of genotype: *F*(1,30)=17.396, *p*<0.001, partial η^2^=0.367; Figure 2A; with no other significant main or interaction effects (all p’s>0.657). Concentrations of tau and p-tau in the cortex were also analyzed, but there were no significant main or interaction effects (all p’s>0.552; Table 1). Primiparous rats were more likely to display regular estrous cycling than nulliparous rats (*χ^2^*=9.992, *p*=0.002; Figure 2B), regardless of genotype (effect of genotype: *p*=0.184). Maternal care behavior, body weight, and adrenal to body weight ratio were not significantly affected by primiparity or hAPOEε4 genotype (all p’s>0.103; see Supplement Table 1).

**Figure 2.**
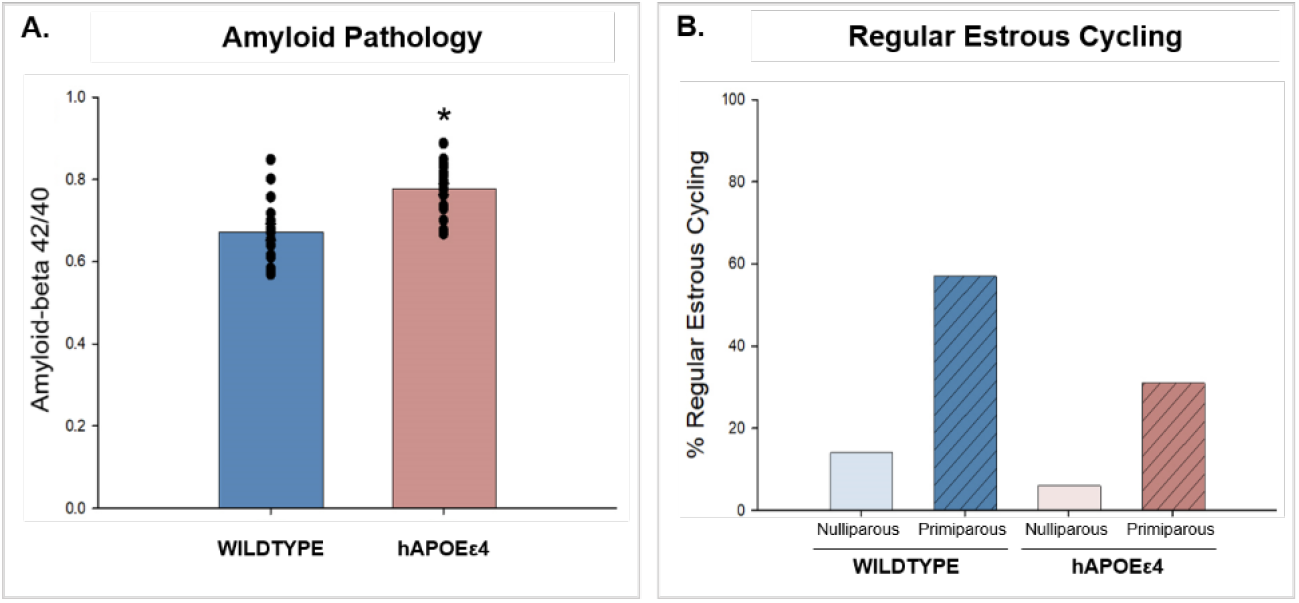
(A) Mean Aβ42 to Aβ40 concentration ratio in the cortex. hAPOEε4 rats showed a higher ratio, which indicates greater amyloid pathology, compared to wildtype rats. (B) Percentage of animals that displayed regular estrous cycling across the experiment. Primiparous rats were more likely to show regular estrous cycling compared to nulliparous rats. * indicates p<0.05. Aβ – amyloid-beta, hAPOEε4 – humanized APOEε4

### 3.2. hAPOEε4 rats made more errors than wildtype rats during initial training of the radial arm maze task. Nulliparous hAPOEε4 rats made more consecutive adjacent arm entries than wildtype rats, and primiparous hAPOEε4 rats made increasingly more consecutive adjacent arm entries

As expected, hAPOEε4 rats made more across-phase errors and a trend for more within-phase errors than wildtype rats (across-phase errors main effect of genotype: *F*(1,41)=4.886, *p*=0.033, partial η^2^=0.106; within-phase errors main effect of genotype: p=0.085; Figure 3A and B). Furthermore, errors decreased across time, although for within phase errors, this approached significance (across-phase errors main effect of block*: F*(3,123)=3.923, *p*=0.010, partial η^2^=0.087; within-phase errors main effect of block: *F*(3,123)=2.525, *p*=0.061, partial η^2^=0.058; Figure 3A and B). There were no other significant main or interaction effects on errors (all p’s>0.135).

**Figure 3.**
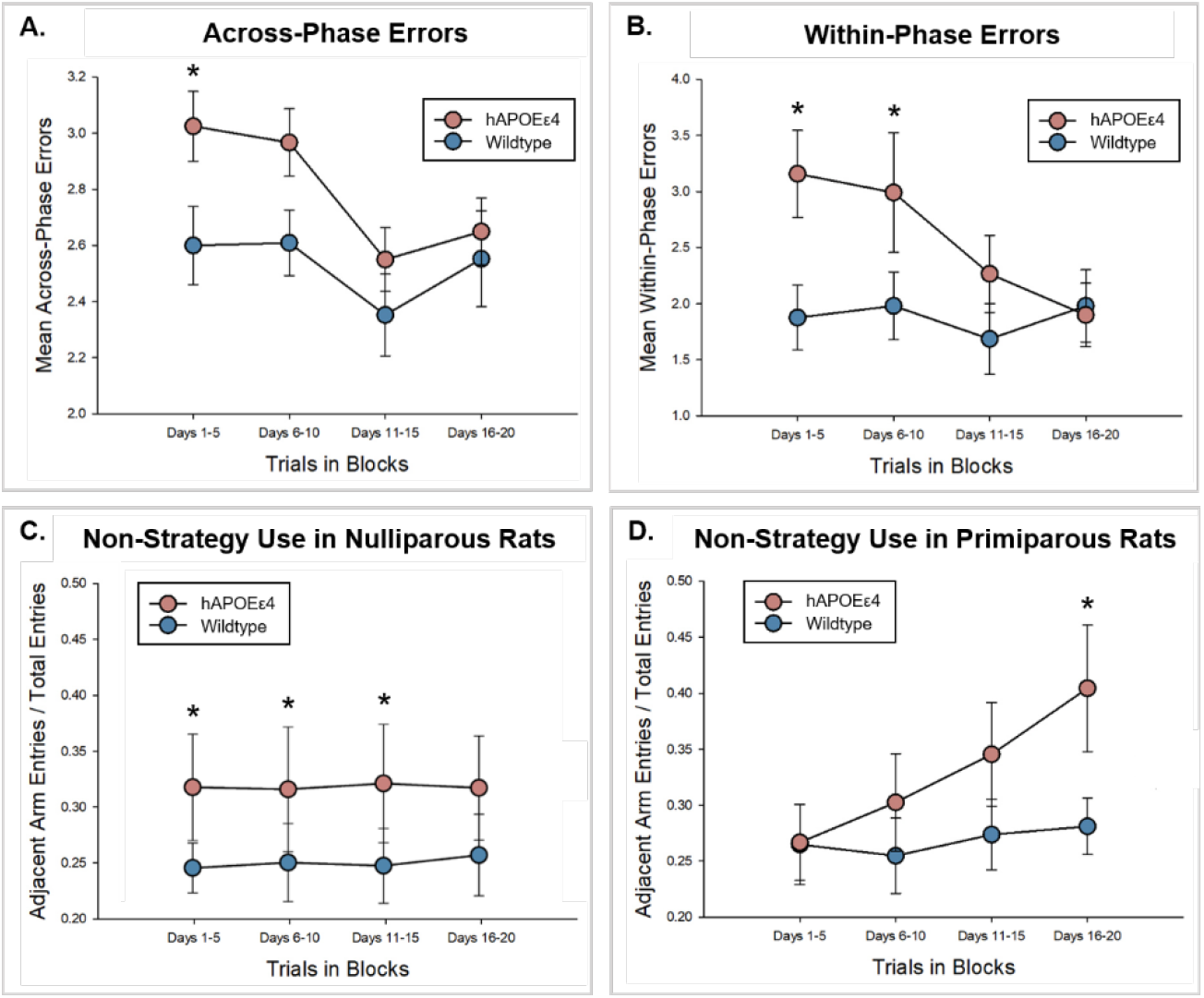
Mean number of across-phase errors (A) and within-phase errors (B) ± standard error of the mean in 5-day blocks in wildtype and hAPOEε4 rats. hAPOEε4 rats made more across-phase errors in block 1 (days 1-5), and more within-phase errors in blocks 1 (days 1-5) and 2 (days 6-10) relative to wildtype rats. Mean ratio of the number of consecutive adjacent arm entries to total entries in 5-day blocks in wildtype and hAPOEε4 (C) nulliparous rats and (D) primiparous rats. More consecutive adjacent arm entries is indicative of non-spatial strategy use. Nulliparous hAPOEε4 made more adjacent arm entries than wildtype rats in blocks 1 (days 1-5), 2 (days 6-10), and 3 (days 11-15). Primiparous hAPOEε4 rats made more adjacent arm entries than wildtype rats in block 4 (days 16-20). * indicates p<0.05 compared to wildtype. hAPOEε4 – humanized APOEε4.

We next examined potential differences in non-spatial strategy use during task acquisition. Relative to the total number of errors, nulliparous hAPOEε4 rats had more consecutive adjacent arm entries than wildtype rats across all blocks except block 4, indicating greater use of a non-spatial strategy (block 1 *p*=0.022, block 2 *p*=0.033, block 3, *p*=0.017, block 4 *p*=0.059 Figure 3C). Primiparous hAPOEε4 rats made more consecutive adjacent arm entries than wildtype rats in blocks 3 and 4 (p’s <0.009) and between all other groups in block 4, indicating greater use of a non-spatial strategy that increased with time (all p’s <0.0002; block by genotype by parity interaction: *F*(3,123)=3.710, *p*=0.013, partial η^2^=0.083; Figure 3D). There was also a significant main effect of block (main effect of block: *F*(3,123)=6.580, *p*<0.001, partial η^2^=0.138) and significant interactions of block by genotype (*F*(3,123)=2.657, *p*=0.051, partial η^2^=0.061) and block by parity (*F*(3,123)=5.030, *p*=0.003, partial η^2^=0.109), but no other significant main or interaction effects on non-spatial strategy use (all p’s>0.124).

### 3.3. Primiparous wildtype rats had more DCX-IR cells, whereas primiparous hAPOEε4 rats had fewer DCX-IR cells, compared to nulliparous rats in the ventral dentate gyrus

To examine potential regional differences within the dentate gyrus, we examined markers in the dorsal and ventral dentate gyrus. Sox2-IR cells were quantified to examine potential differences in the number of neural stem cells in the dentate gyrus. There were significantly more Sox2-IR cells in the ventral dentate gyrus (main effect of region: *F*(1,28)=401.741, *p*<0.001, partial η^2^=0.935; Figures 4A, B) than the dorsal dentate gyrus. There were no other significant main or interaction effects on Sox-IR cells (all p’s > 0.163).

**Figure 4.**
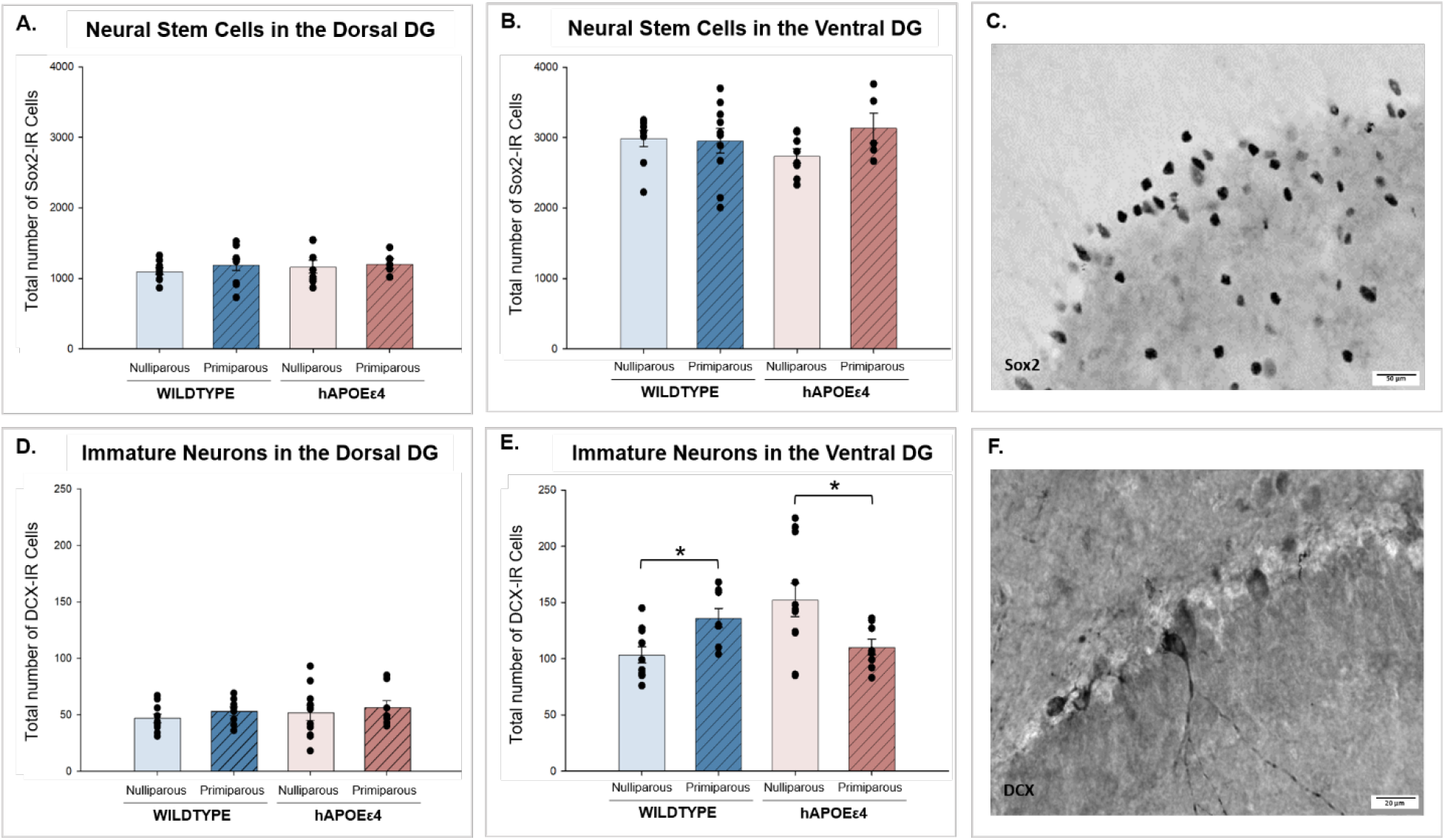
Total number of sampled Sox2-IR cells ± standard error of the mean in the (A) dorsal dentate gyrus and (B) ventral dentate gyrus. There were more Sox2-IR cells in the ventral dentate gyrus than the dorsal dentate gyrus. (C) Photomicrograph of Sox2-IR cells. Total number of DCX-IR cells in the (D) dorsal and (E) ventral dentate gyrus. In the ventral dentate gyrus, primiparous wildtype rats had more immature neurons than nulliparous wildtype rats, and primiparous hAPOEε4 rats had fewer immature neurons than nulliparous hAPOEε4 rats. Nulliparous hAPOEε4 rats also had more immature neurons than wildtype nulliparous rats. (F) Photomicrograph of DCX-IR cells. * indicates p<0.05. Sox2 – SRY-box transcription factor 2, DCX – doublecortin, DG – dentate gyrus, hAPOEε4 – humanized APOEε4.

DCX-IR cells were quantified to examine potential differences in the number of immature neurons in the dentate gyrus. Among the wildtype group, primiparous rats had significantly more DCX-IR cells in the ventral dentate gyrus than nulliparous rats (*p*=0.050, Cohen’s *d*=0.528; region by genotype by parity interaction: *F*(1,33)=8.096, *p*=0.008 partial η^2^=0.197), whereas among the hAPOEε4 group, primiparous rats had significantly fewer DCX-IR cells in the ventral dentate gyrus than nulliparous rats (*p*=.007, Cohen’s *d*=0.220; Figure 4E). Furthermore, nulliparous hAPOEε4 rats had significantly more DCX-IR cells than nulliparous wildtype rats (*p*=0.003, Cohen’s *d*=0.275). However, there were no significant differences in the number of DCX-IR cells with parity and genotype in the dorsal dentate gyrus (p’s>0.809). There was a significant main effect of region (*F*(1,33)=132.084, *p*<0.001, partial η^2^=0.800; Figures 4D, E) but no other significant main or interaction effects on DCX-IR cells (all p’s>0.193).

### 3.4. Primiparous hAPOEε4 rats had the lowest percentage of DCX/zif268 co-labelled cells compared to all other groups in the dorsal and ventral dentate gyrus

DCX/zif268 co-labelled cells were quantified to examine potential differences in activation of immature neurons in the dentate gyrus. Primiparous hAPOEε4 rats had a lower percentage of DCX/zif268 co-labelled cells compared to all other groups (genotype by parity interaction: *F*(1,36)=10.186, *p*=0.003, partial η^2^=0.221; Figure 5A,B). There was a significant region by parity interaction, where primiparous rats had a lower percentage of DCX/zif268 co-labelled cells compared to nulliparous rats in the dorsal dentate gyrus F(1,36)=11.266, *p*=0.002, partial η^2^=0.238; Figure 5A), and planned comparisons revealed that this effect was stronger in the hAPOEε4 group (hAPOEε4: *p*<0.001, Cohen’s *d*=2.256; wildtype: *p*=0.053, Cohen’s *d*=0.192. There were also significant main effects of parity (*F*(1,36)=6.407, *p*=0.016, partial η^2^=0.151), and region (*F*(1,36)=113.445, *p*<0.001, partial η^2^=0.759), but no other significant main effects or interactions (all p’s>0.083).

**Figure 5.**
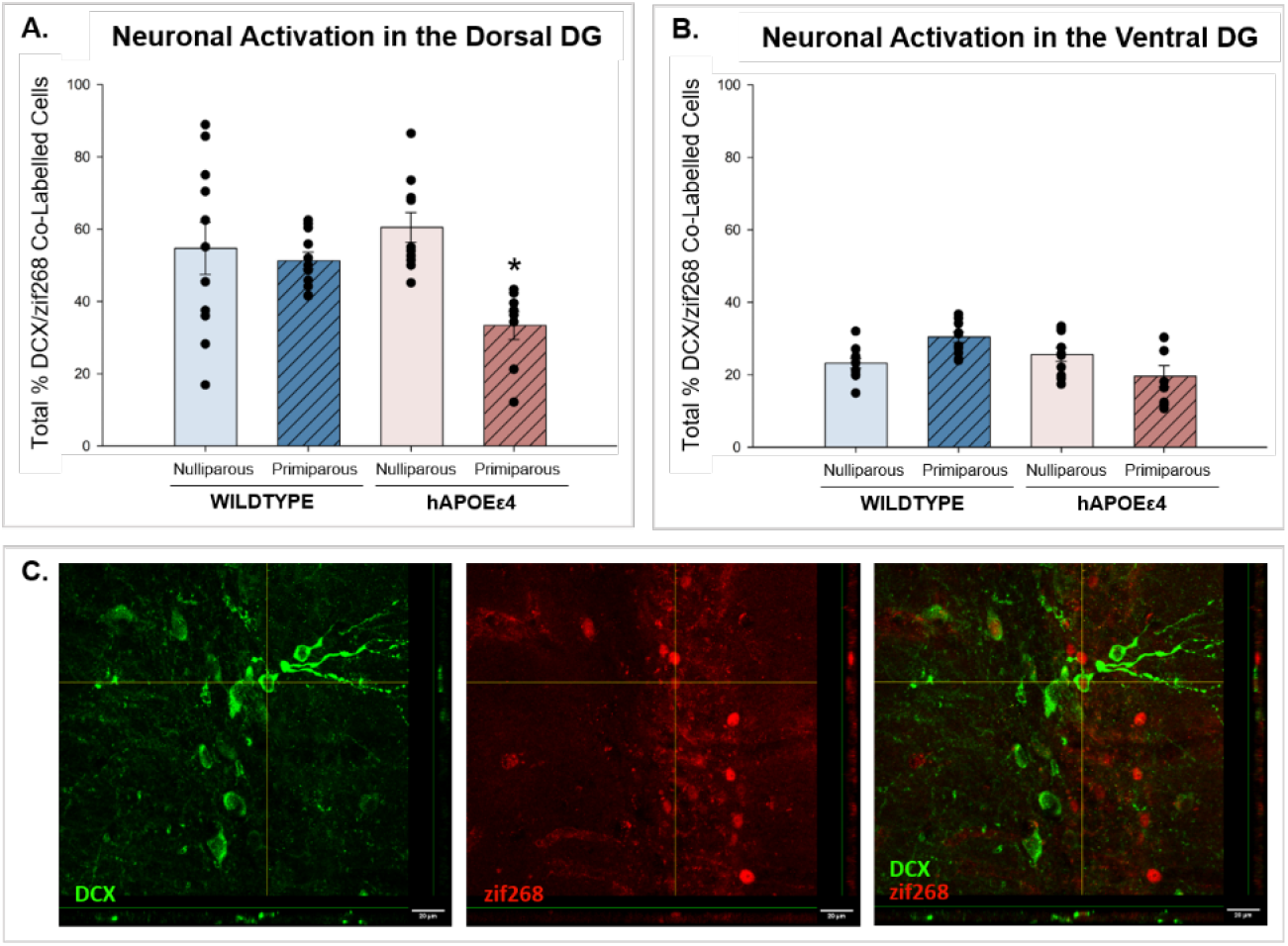
Total percentage of DCX/zif268 co-labelled cells ± standard error of the mean in the (A) dorsal dentate gyrus and (B) ventral dentate gyrus. There was a lower percentage of DCX/zif268 co-labelled cells in primiparous hAPOEε4 rats compared to all other groups. (C) Photomicrographs of a DCX and zif268 co-labelled cell. * indicates p<0.05 compared to all other groups. DCX – doublecortin, zif268 – zinc finger-containing transcription factor 268, DG – dentate gyrus, hAPOEε4 – humanized APOEε4.

### 3.5. hAPOEε4 rats had fewer Iba1-IR cells than wildtype rats in the ventral dentate gyrus

Iba1-IR cells were quantified to examine potential differences in the number of microglia in the dentate gyrus. hAPOEε4 rats had fewer Iba1-IR cells than wildtype rats in the ventral, but not dorsal, region (region by genotype interaction: *F*(1,28)=4.056, *p*=0.050, partial η^2^=0.127; Figure 6B). There was a significant main effect of region (*F*(1,28)=90.396, *p*<0.001, partial η^2^=0.764; Figures 6A, B), but no other significant main effects or interactions (all p’s>0.155).

**Figure 6.**
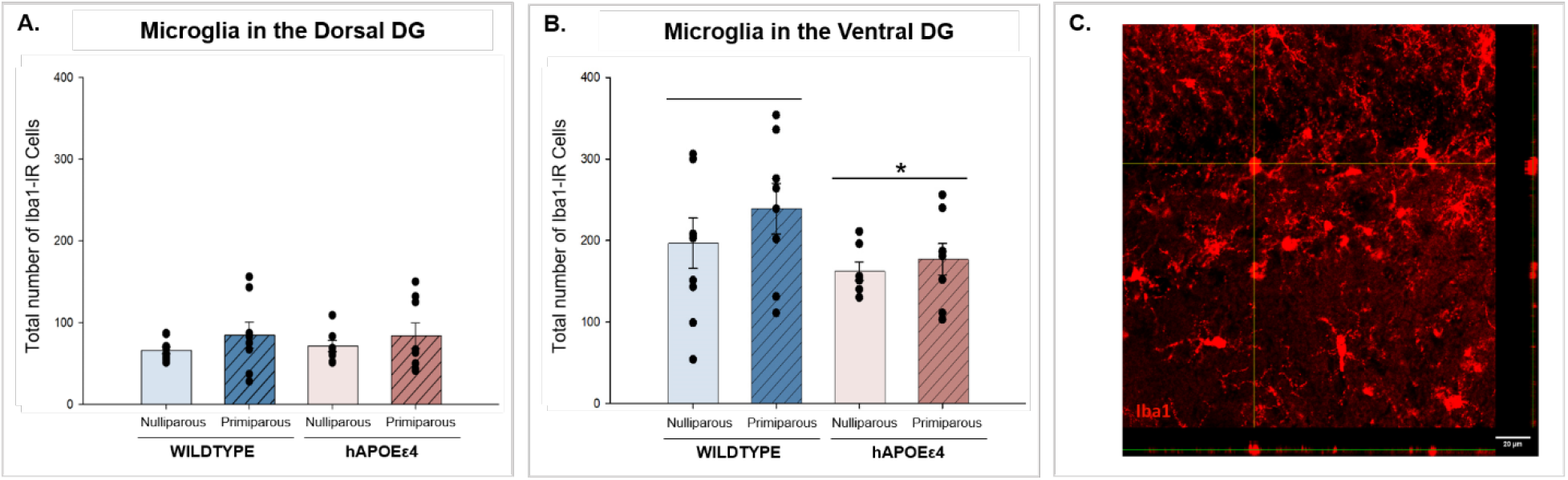
Total number of Iba1-IR cells ± standard error of the mean in the (A) dorsal dentate gyrus and (B) ventral dentate gyrus. There were more Iba1-IR cells in the ventral dentate gyrus than the dorsal dentate gyrus. In the ventral dentate gyrus, hAPOEε4 rats had fewer Iba1-IR cells than wildtype rats. (C) Photomicrograph of Iba1-IR cells. * indicates p<0.05. Iba1 – ionized calcium-binding adaptor molecule 1, DG – dentate gyrus, hAPOEε4 – humanized APOEε4.

### 3.6. Expression of IL-1β, IL-4, and IL-5 were highest in the dorsal hippocampus. Expression of CXCL1 was higher in the dorsal and ventral hippocampus compared to the frontal cortex

ANOVAs were conducted for each of the 8 cytokines and 1 chemokine detected in the assay (see Supplement Figure 1). The dorsal hippocampus showed the highest expression of IL-1β (compared to ventral hippocampus: *p*=0.020; compared to frontal cortex: *p*=0.004; main effect of region: *F*(2,58)=5.378, *p*=0.007, partial η^2^=0.156), IL-4 (compared to ventral hippocampus: *p*=0.004; compared to frontal cortex: *p*=0.005; main effect of region: *F*(2,58)=5.860, *p*=0.005, partial η^2^=0.168), and IL-5 (compared to ventral hippocampus: *p*=0.001; compared to frontal cortex: *p*=0.0005; main effect of region: *F*(2,46)=9.473, *p*<0.001, partial η^2^=0.292). Expression of CXCL1 was higher in the dorsal hippocampus (*p*=0.001) and ventral hippocampus (*p*=0.005) compared to the frontal cortex (main effect of region: *F*(2,50)=7.203, *p*=0.002, partial η^2^=0.224). There were no other significant main or interaction effects involving parity or genotype for expression of IL-1β (all p’s>0.070), IL-5 (all p’s>0.082), IL-4 (all p’s>0.077), CXCL1 (all p’s>0.122), IL-10 (all p’s>0.120), IFN-γ (all p’s>0.096), IL-6 (all p’s>0.246), and TNF-α (all p’s>0.333) in the dorsal hippocampus, ventral hippocampus, and frontal cortex.

#### 3.6.1 Primiparous hAPOEε4 rats a higher Th1/Th2 ratio in the ventral hippocampus than primiparous wildtype rats. Primiparous rats had a higher Th1/Th2 ratio in the frontal cortex than nulliparous rats

Th1 to Th2 z-score ratios were analyzed to examine the balance between Th1-related cytokines (IL-1β, IFN-γ, TNF-α) and Th2-related cytokines (IL-4, IL-5, IL-10, IL-13). There were no significant main or interaction effects within the dorsal hippocampus (all p’s>0.267; Figure 7A). In the ventral hippocampus, primiparous hAPOEε4 rats had higher Th1/Th2 than primiparous wildtype rats (genotype by parity interaction: *F*(1,39)=6.010, *p*=0.019, partial η^2^=0.134; Figure 7B). There were no significant main effects in the ventral hippocampus (all p’s>0.185). In the frontal cortex, primiparous rats had higher Th1/Th2 than nulliparous rats (main effect of parity: *F*(1,39)=5.106, *p*=0.030, partial η^2^=0.116; Figure 7C). There were no other significant main or interaction effects in the frontal cortex (all p’s>0.176).

**Figure 7.**
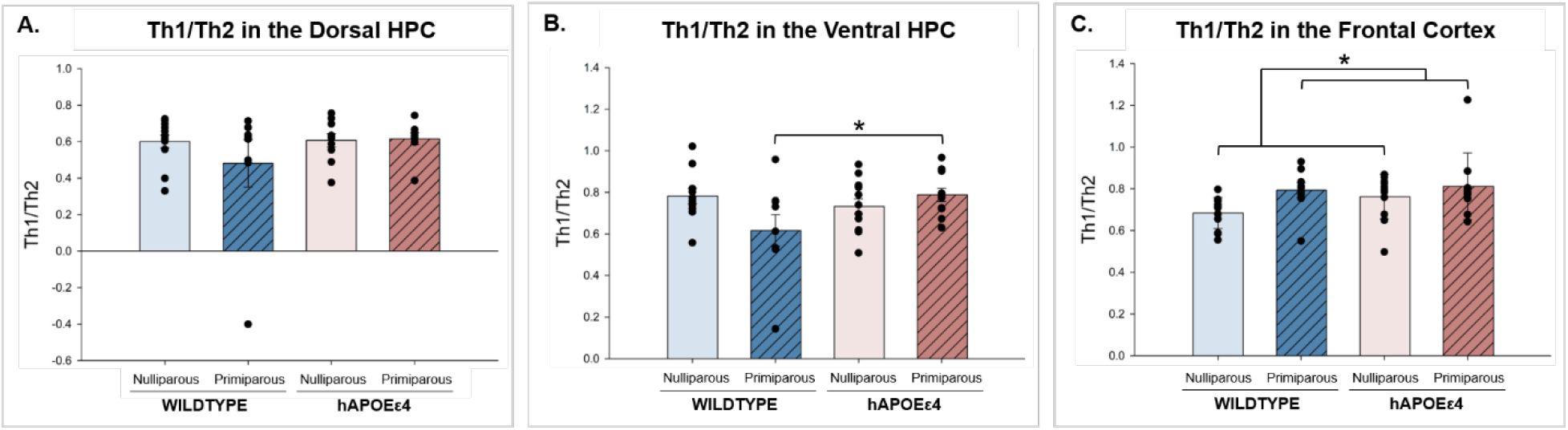
Th1 to Th2 ratio ± standard error of the mean in the (A) dorsal hippocampus, (B) ventral hippocampus, and (C) frontal cortex. In the ventral hippocampus, primiparous hAPOEε4 rats had higher Th1/Th2 than wildtype primiparous rats. In the frontal cortex, primiparous rats had higher Th1/Th2 than nulliparous rats. * indicates p<0.05. Th–T helper, HPC – hippocampus, hAPOEε4 – humanized APOEε4.

### 3.7. hAPOEε4 rats had higher concentrations of tryptophan metabolites (kynurenine, kynurenic acid, and anthranilic acid) than wildtype rats. Primiparous rats had a greater kynurenine to tryptophan ratio than nulliparous rats

Plasma concentrations of kynurenine, kynurenic acid, and anthranilic acid were significantly higher in hAPOEε4 compared to wildtype rats (main effect of genotype: kynurenine: *F*(1,39)=1.572, *p*=0.004, partial η^2^=0.189, kynurenic acid*: F*(1,40)=21.639, *p*<0.001, partial η^2^=0.351; anthranilic acid: *F*(1,38)=13.329, *p*=0.001, partial η^2^=0.260; Figures 8A,B,C). Kynurenine concentrations were also significantly higher in primiparous than nulliparous rats (main effect of parity: *F*(1,39)=16.635, *p*<0.001, partial η^2^=0.299; Figure 8A). Planned comparisons revealed significantly higher anthranilic acid concentration in hAPOEε4 primiparous than wildtype primiparous rats (*p*=0.006, Cohen’s *d*=1.235). There were no other significant main or interaction effects for these or any other metabolites (all p’s>0.165). Other metabolites that were included in the assay also showed no significant main or interaction effects.

**Figure 8.**
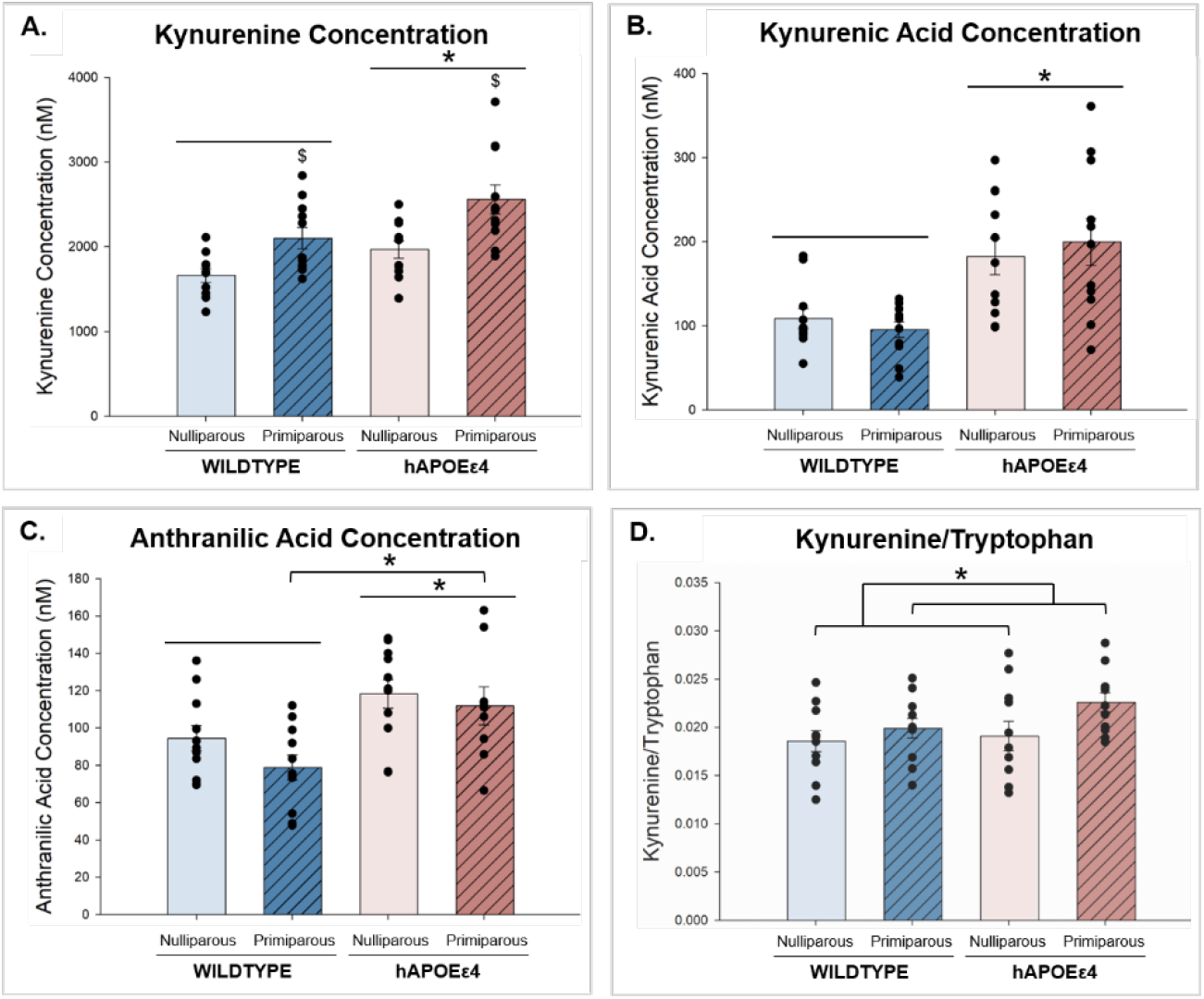
(A) Plasma concentrations of kynurenine ± standard error of the mean. hAPOEε4 rats had greater levels of kynurenine than wildtype rats. $ indicates that primiparous rats had greater levels of kynurenine than nulliparous rats. (B) Plasma concentrations of kynurenic acid. hAPOEε4 rats had greater levels of kynurenic acid than wildtype rats. (C) Plasma concentrations of anthranilic acid. hAPOEε4 rats had greater levels of anthranilic acid than wildtype rats and hAPOEε4 primiparous rats had greater levels of anthranilic acid than wildtype primiparous rats. (D). Kynurenine to tryptophan ratio. Primiparous rats had higher kynurenine/tryptophan than nulliparous rats. * indicates p<0.05. hAPOEε4 – humanized APOEε4.

To further investigate potential differences in tryptophan metabolism, kynurenine to tryptophan ratios were analyzed. Primiparous rats had significantly higher kynurenine/tryptophan than nulliparous rats (main effect of parity: *F*(1,40)=4.230, *p*=0.046, partial η^2^=0.096; Figure 8D). This effect seems to be driven by the hAPOEε4 group, although it was not significant after Bonferroni correction (*p*=0.041, Cohen’s *d*=0.819). There were no other main or interaction effects for kynurenine/tryptophan ratio (all p’s>0.180).

## 4. Discussion

In the present study, a history of parity influenced cognitive and brain-based biomarkers at middle age in a manner that depended on hAPOEε4 genotype. Primiparous hAPOEε4 rats increased use of a non-spatial strategy and had fewer and less active new-born immature neurons in the dentate gyrus compared to all other groups. hAPOEε4 genotype, regardless of parity, was associated with shifts in neuroinflammatory markers and tryptophan metabolism. In the ventral hippocampus, hAPOEε4 genotype reduced the number of microglia, yet primiparous hAPOEε4 rats exhibited higher Th1/Th2, indicating a bias towards a pro-inflammatory phenotype, compared to wildtype rats. An assessment of tryptophan metabolomics revealed that hAPOEε4 rats had higher plasma concentrations of tryptophan metabolites than wildtype rats, and parity was associated with greater activation of the kynurenine pathway. Together, these data demonstrate that primiparous hAPOEε4 rats elicit use of a non-spatial strategy, which is accompanied with decreased recruitment of new neurons and increased pro-inflammation in the hippocampus.

### 4.1. hAPOEε4 rats made more spatial memory errors and failed to use a spatial strategy in the radial arm maze compared to wildtype rats. Primiparous hAPOEε4 rats increased use of a non-spatial strategy in the radial arm maze

hAPOEε4 rats made more errors in the spatial working memory task than wildtype rats. Performance on this task relies on the integrity of both the hippocampus and the frontal cortex, suggesting that hAPOEε4 rats may experience impairments in either of these regions or the connectivity between these regions (Floresco et al., 1997). This aligns with literature that demonstrate functional connectivity disruptions between the hippocampus and the frontal lobes in individuals with AD (Allen et al., 2007; Josephs et al., 2017). Among nulliparous rats, the hAPOEε4 group tended to use a non-spatial strategy, implicating decreased recruitment of the hippocampus during the task, more than the wildtype group. This is consistent with studies in humans showing decreased activation of the hippocampus with aging in APOEε4 carriers (Håglin et al., 2023). Importantly, the present study is the first to investigate and show that parity further impacts this relationship, as primiparous hAPOEε4 rats were the only group to increase the use of a non-spatial strategy to solve the task. Together, this data supports literature showing declines in spatial strategy use with both aging and AD, which leads to increased recruitment of compensatory navigation strategies (Colombo et al., 2017; Parizkova et al., 2018; Wiener et al., 2013). In particular, as AD advances, there is increased preference for an egocentric over an allocentric strategy (Parizkova et al., 2018). This compensatory mechanism could be a possible explanation for hAPOEε4 rats adopting a non-spatial strategy that is more egocentric based, which becomes increasingly prominent throughout training in the primiparous group.

In this study, previous parity did not influence the number of errors made in the radial arm maze, which is inconsistent with previous findings using tasks that rely on the integrity of the hippocampus. Rodent studies have generally reported higher spatial working and reference memory scores in early acquisition of hippocampus-dependent tasks with parity, in middle age (Barha et al., 2015; Cui et al., 2014; Galea et al., 2018). As mentioned, performance in the cognitive task used in the present study relies on both the hippocampus and frontal cortex, implicating that perhaps these two areas may be differentially affected by parity, resulting in no overt differences in the number of errors made in the task. In fact, we also find region-specific effects of parity and hAPOEε4 on neuroinflammation that may contribute to these cognitive findings. In sum, findings from the present study are suggestive that parity may differentially influence strategy use, reflecting decreased recruitment of the hippocampus, depending on hAPOEε4 genotype.

### 4.2. Previous primiparity increased neurogenesis in wildtype rats but reduced it in hAPOEε4 rats in the **ventral dentate gyrus.**

Primiparous wildtype rats had greater levels of neurogenesis than nulliparous wildtype rats in the ventral dentate gyrus, similar to previous work showing greater levels of neurogenesis in middle-aged primiparous and multiparous rats compared to nulliparous rats (Barha et al., 2015; Eid et al., 2019). This finding also aligns with human brain imaging data from the UK Biobank that shows evidence of less brain aging in middle-aged parous females compared to nulliparous females (de Lange et al., 2019). Together, these data demonstrate that parity contributes long-lasting effects on neuroplasticity and underscore the importance of considering female-unique experiences like parity in research on brain aging.

In contrast to what is seen in wildtype rats, primiparous hAPOEε4 rats had fewer immature neurons compared to nulliparous hAPOEε4 rats. This may be explained by the healthy cell bias, which was first proposed by Brinton (2008) to explain that as neurological health progresses from healthy to unhealthy, the actions of estrogens may also progress from protective to detrimental. A recent study using an APOE-TR^+/+^/5xFAD^+/-^ mouse model found that the neuroprotective effects of 17β-estradiol on hippocampal memory and plasticity were seen in mice with no (E3FAD) or one (E3/E4FAD) hAPOEε4 alleles, but impeded in mice with both (E4/E4FAD) alleles (Taxier et al., 2022). Furthermore, Cui et al. (2014) show that parity is associated with a shorter latency to solve a spatial working memory task in wildtype mice, but a longer latency to solve the task in APP23 mice. Together, this implicates that parity can exert either protective or detrimental actions on brain aging, depending on AD risk, including APOEε4 genotype. Indeed, the findings from the present study illustrate that although primiparous wildtype rats show increased neurogenesis at middle age, primiparous hAPOEε4 parous rats show decreased neurogenesis at middle age.

In the present study, we found that among nulliparous rats, there were more immature neurons in the hAPOEε4 group relative to the wildtype group, and this is consistent with other studies showing increased hippocampal neurogenesis in early AD in both rodent models (Adeosun et al., 2019; Jin, Galvan, et al., 2004; Koutseff et al., 2014) and humans (Jin, Peel, et al., 2004; Lee et al., 2023). Together, these findings support the possibility of a compensatory mechanism to replace damaged neurons from AD-related neuropathology. Conversely, it is also possible that females show a different trajectory of neurogenesis with APOEε4 genotype or that aging impacts the integrity of the hippocampus uniquely in APOEε4 females – future studies are needed to evaluate these possibilities. Given that the rats were examined at 13-14 months old, we expect that at an older age, we may in turn see a decline in neurogenesis. Lastly, there were no genotype or parity differences in the number of neural stem cells in the dentate gyrus of the hippocampus. As such, our findings demonstrate that despite having similar numbers of neural stem cells, parity differentially affected the number of new-born neurons in wildtype and hAPOEε4 rats. More research is needed to investigate the progression of new-born immature neurons to elucidate whether hAPOEε4 genotype further impacts maturation, survival, and integration.

### 4.3. Primiparous hAPOEε4 rats had less activation of new-born neurons compared to primiparous wildtype rats in the dorsal and ventral dentate gyrus

hAPOEε4 primiparous rats had less activation of immature neurons compared to wildtype primiparous rats in the dentate gyrus in response to memory retrieval, which parallels our finding that hAPOEε4 primiparous rats increased non-spatial strategy use to solve the radial arm maze. Together, this indicates that zif268 activation may be necessary for spatial strategy use. In humans, APOEε4 carriers show decreased hippocampal activation with aging, whereas non-carriers show no differences (Håglin et al., 2023), although sex had not been examined. Despite the differences in zif268 activation, there were no differences in number of errors across groups, indicating that hAPOEε4 primiparous rats may be relying on different features of the hippocampus or frontal cortex to engage in spatial working memory. Further research is needed to identify the mechanisms underlying the expression of different IEGs and other regions in response to cognitive tasks depending on factors like parity and hAPOEε4 genotype.

### 4.4. hAPOEε4 rats had fewer microglia in the ventral dentate gyrus compared to wildtype rats

In the present study, we showed that hAPOEε4 rats had fewer microglia than wildtype rats in the ventral dentate gyrus. The role of microglia in the pathogenesis of AD is complex, as microglia have been implicated in both neuroprotective and neurotoxic ways (Kinney et al., 2018; Streit, 2005). One hypothesis is that presence of Aβ drives activation of microglia to phagocytose and clear Aβ from the brain, however, sustained activation of microglia can lead to accumulation of Aβ and exacerbation of AD pathology (Baik et al., 2016; Bolmont et al., 2008; Stalder et al., 1999). However, majority of studies in this area have focused on males, resulting in a gap in our understanding about neuroinflammatory phenomena in AD, in females. Intriguingly, one study using a model of vascular contributions to cognitive impairment and dementia reported an increase in the number of hippocampal microglia in male, but not female middle-aged mice (Abi-Ghanem et al., 2023). This is partly consistent with our findings – and although Abi-Ghanem et al. (2023) did not find a decrease in microglia in females, it is important to consider that the model of dementia used is different from that of the present study and involves compromised cardiovascular and cerebrovascular health, which have unique implications for brain health and inflammation. Nonetheless, the findings from the present study and that of Abi-Ghanem et al. (2023) support the idea that there may be distinct neuroimmune pathways or mechanisms in AD between males and females, and these largely remain to be investigated. Indeed, Casaletto et al. (2022) found striking sex differences in the pathways through which microglia may contribute to AD neuropathology. They show a bidirectional relationship between Aβ and microglial activation that accounts for tau burden in females, and direct independent relationships between microglial activation or Aβ with tau in males. As mentioned earlier, we did not observe any genotype differences with tau burden. This is possibly in line with the work by Casaletto et al. (2022), as we found low levels of microglia and no detectable increases in tau in our hAPOEε4 female rats. More research is needed to disentangle the relationship between neuroinflammatory markers like microglia and AD-related pathology in females.

Lastly, the present study did not find any effect of parity on the number of microglia in the dentate gyrus, which is consistent with previous research in primiparous rats (Duarte-Guterman et al., 2023; Eid et al., 2019).

Intriguingly, the effects of hAPOEε4 genotype and parity on neurogenesis (neural stem cells and immature new-born neurons) and microglia were only significant in the ventral sub-region of the dentate gyrus. The dorsal and ventral hippocampus have long been shown to possess distinct functional capabilities, with the dorsal hippocampus move involved in learning and memory associated with navigation, exploration, and locomotion, and the ventral hippocampus more involved in motivational and emotional behaviour, including stress (Fanselow & Dong, 2010). Although both the dorsal and ventral hippocampus are important for efficient spatial navigation, spatial working memory acquisition and retrieval rely more heavily on the ventral hippocampus whereas spatial reference memory performance is modulated by the dorsal hippocampus (Moser & Moser, 1998). Thus, it is perhaps not surprising that the effects we found were predominantly in the ventral dentate gyrus.

### 4.5. Primiparous hAPOEε4 rats had a greater pro-inflammatory phenotype in the ventral hippocampus compared to wildtype rats. In the frontal cortex, primiparous rats had a greater pro-inflammatory phenotype compared to nulliparous rats, regardless of hAPOEε4 genotype

We investigated the balance between Th1- and Th2-related cytokines and found that in the ventral hippocampus, primiparous hAPOEε4 rats had higher Th1/Th2 compared to wildtype rats, suggesting dominance of Th1-related immunity. Although both Th1 and Th2 cells are essential for the maintenance of a healthy milieu within the central nervous system, an imbalance in the ratio between Th1 and Th2 cells has been shown to contribute to neurodegeneration (Luckheeram et al., 2012). Indeed, Th1 cells directly contribute to neuroinflammation through the release of pro-inflammatory cytokines and have been linked to impaired cognitive function and elevated Aβ deposition (Browne et al., 2013). In the frontal cortex, primiparous rats had higher Th1/Th2 compared to nulliparous rats regardless of hAPOEε4 genotype. Together, these findings reinforce that parity and hAPOEε4 genotype influence the neuroinflammatory milieu in a region-dependent manner.

### 4.6. hAPOEε4 rats showed greater tryptophan metabolism than wildtype rats. Primiparous rats showed greater activation of the kynurenine pathway than nulliparous rats

We report that hAPOEε4 rats had higher plasma concentrations of the tryptophan metabolites kynurenine, kynurenic acid, and anthranilic acid, compared to wildtype rats. These findings are consistent with other studies showing increased activation of the kynurenine pathway in AD patients and animal models (Castro-Portuguez & Sutphin, 2020; Chatterjee et al., 2018; W. Wu et al., 2013).

There were no effects of parity on kynurenine pathway metabolite concentrations, consistent with other work (Duarte-Guterman et al., 2023). However, calculating the kynurenine to tryptophan ratio allowed us to directly measure the state of activation of the kynurenine pathway, which we did find to be increased in primiparous rats, with a trend towards a greater effect in the hAPOEε4 group. Enzymatic activity along the pathway influences either or both kynurenine and tryptophan and are thus the primary determinants of this ratio. In particular, this ratio is frequently examined as an indicator of inflammation-mediated tryptophan metabolism via the enzyme indoleamine 2,3-dioxygenase in the pathway (Oxenkrug, 2010). Together, these data implicate the kynurenine pathway as a potential mediator of the effects of hAPOEε4 genotype on spatial working memory (more errors) and neuroinflammation in the ventral hippocampus (lower levels of microglia). Continued research in this area will be needed to elucidate the mechanisms underlying these relationships.

## 5. Conclusions

We found that in middle-aged rats, previous primiparity had opposing effects on spatial strategy use and neuroplasticity, depending on hAPOEε4 genotype. This research showed that primiparous hAPOEε4 rats increased use of a non-spatial strategy and had fewer and less active new-born immature neurons in the dentate gyrus, suggesting recruitment of different features of the hippocampus or frontal cortex during the cognitive task in these rats. Indeed, this research also demonstrates that cytokine expression in the hippocampus and frontal cortex are differentially affected by parity and hAPOEε4 genotype. Given that parity has been implicated in AD risk and pathology, it is crucial to understand its long-term influences on the aging brain with hAPOEε4 genotype. Collectively, this research illustrates that although previous parity may be neuroprotective in a healthy state, it may further impair AD-related biomarkers in the brain, including neuroplasticity and neuroinflammation, depending on APOEε4 genotype. This work underscores the importance of considering reproductive history and hAPOEε4 genotype in aging and AD research.

## 6. Supplementary Material

**Table S1.**
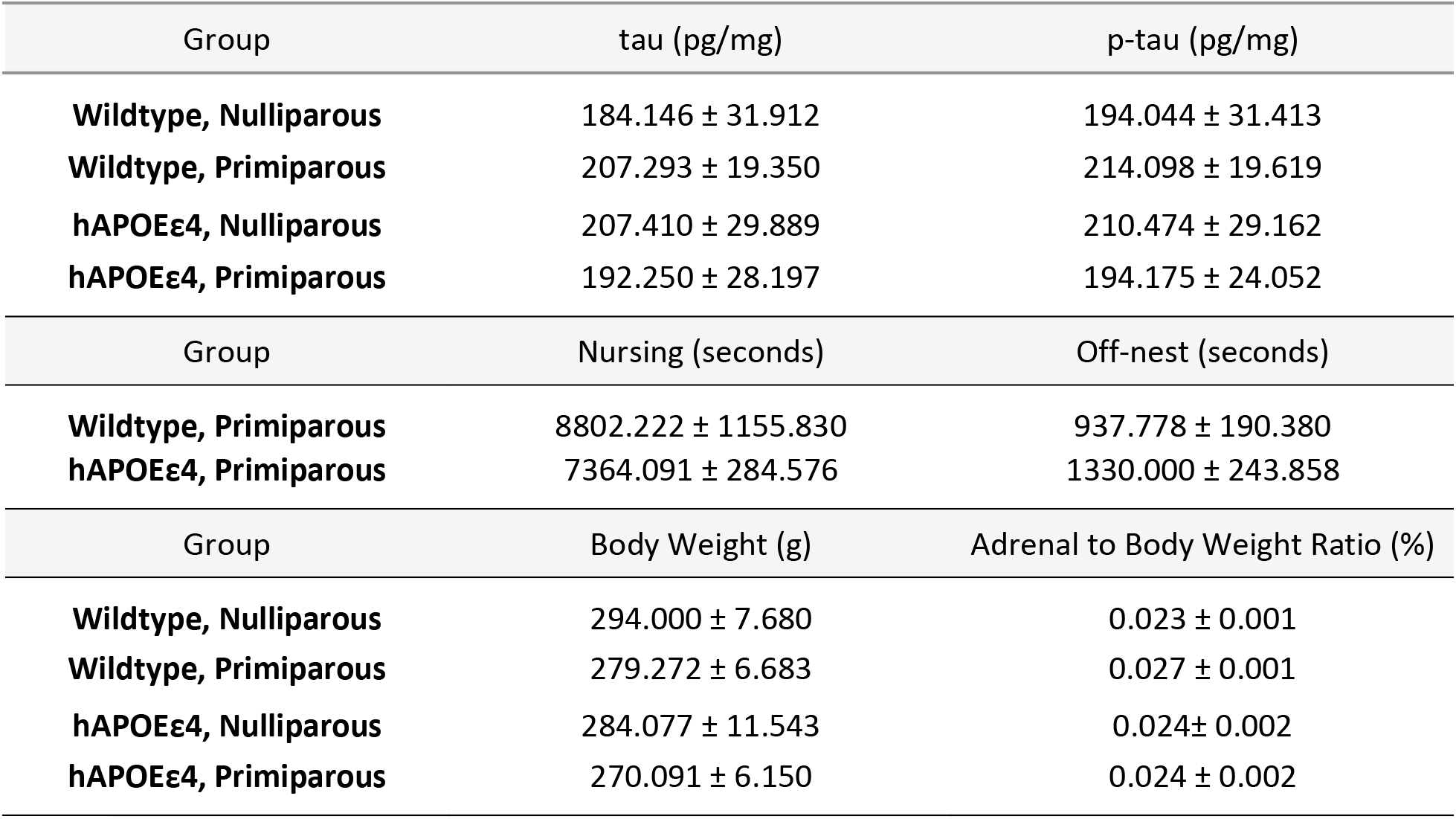
Mean concentrations of tau and p-tau, mean time spent nursing (including arched-back, blanket, and passive nursing) and off-nest (including self-grooming and tail chasing), and mean body weight and adrenal to body weight ratio ± standard error of the mean. p-tau – phosphorylated tau, hAPOEε4 – humanized APOEε4.

**Figure S1.**
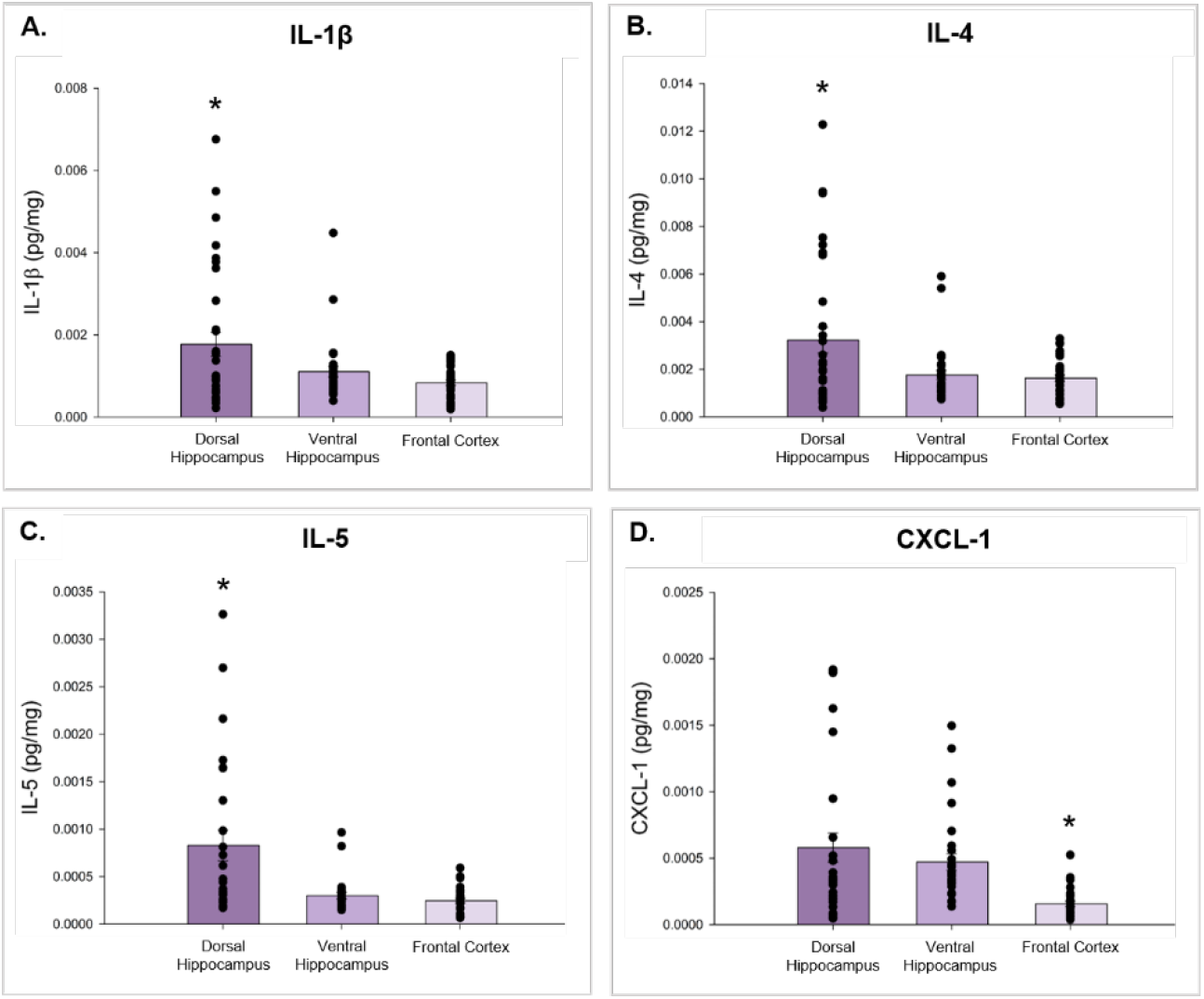
Concentrations ± standard error of the mean of (A) IL-1β, (B) IL-4, (C) IL-5, and (D) CXCL-1 in the dorsal hippocampus, ventral hippocampus, and frontal cortex. Expression of IL-1, IL-4, and IL-5 was greatest in the dorsal hippocampus (compared to ventral hippocampus and frontal cortex), whereas expression of CXCL-1 was greater in the dorsal and ventral hippocampus compared to the frontal cortex. * indicates p<0.05 compared to all other groups. IL-1β–interleukin-1-beta, IL-4–interleukin-4, IL-5– interleukin-5, CXCL1–chemokine (CXC-motif) ligand 1; hAPOEε4 – humanized APOEε4.

## Acknowledgements

We gratefully acknowledge Dr. Stan Floresco for assisting in analysis of the cognitive data and Jared E.J. Splinter for developing and Nadine Truter for optimizing the MATLAB code used for automatic cell counting. We would also like to thank Daria Tai and Nicole Minielly for their help with breeding and animal care as well as the animal care staff at the Centre for Disease Modelling at the University of British Columbia.

## References

1. Abi-Ghanem, C., Salinero, A. E., Kordit, D., Mansour, F. M., Kelly, R. D., Venkataganesh, H., Kyaw, N.-R., Gannon, O. J., Riccio, D., Fredman, G., Poitelon, Y., Belin, S., Kopec, A. M., Robison, L. S., & Zuloaga, K. L. (2023). Sex differences in the effects of high fat diet on underlying neuropathology in a mouse model of VCID. Biology of Sex Differences, 14, 31. https://doi.org/10.1186/s13293-023-00513-y

2. Adeosun, S. O., Hou, X., Shi, L., Stockmeier, C. A., Zheng, B., Raffai, R. L., Weisgraber, K. H., Mosley, T. H., & Wang, J. M. (2019). Female Mice with Apolipoprotein E4 Domain Interaction Demonstrated Impairments in Spatial Learning and Memory Performance and Disruption of Hippocampal Cyto-Architecture. Neurobiology of Learning and Memory, 161, 106–114. https://doi.org/10.1016/j.nlm.2019.03.012

3. Allen, G., Barnard, H., Mccoll, R., Hester, A., Fields, J., Weiner, M., Ringe, W., Lipton, A., Brooker, M., McDonald, E., Rubin, C., & Cullum, M. (2007). Reduced Hippocampal Functional Connectivity in Alzheimer Disease. Archives of Neurology, 64, 1482–1487. https://doi.org/10.1001/archneur.64.10.1482

4. Altmann, A., Tian, L., Henderson, V. W., & Greicius, M. D. (2014). Sex Modifies the APOE-Related Risk of Developing Alzheimer’s Disease. Annals of Neurology, 75(4), 563–573. https://doi.org/10.1002/ana.24135

5. Alzheimer’s Association. (2023). 2023 Alzheimer’s disease facts and figures. Alzheimer’s & Dementia, n/a(n/a). https://doi.org/10.1002/alz.13016

6. Amador-Arjona, A., Cimadamore, F., Huang, C.-T., Wright, R., Lewis, S., Gage, F. H., & Terskikh, A. V. (2015). SOX2 primes the epigenetic landscape in neural precursors enabling proper gene activation during hippocampal neurogenesis. Proceedings of the National Academy of Sciences, 112(15), E1936–E1945. https://doi.org/10.1073/pnas.1421480112

7. Bae, J. B., Lipnicki, D. M., Han, J. W., Sachdev, P. S., Kim, T. H., Kwak, K. P., Kim, B. J., Kim, S. G., Kim, J. L., Moon, S. W., Park, J. H., Ryu, S.-H., Youn, J. C., Lee, D. Y., Lee, D. W., Lee, S. B., Lee, J. J., Jhoo, J. H., Llibre-Rodriguez, J. J.,…for Cohort Studies of Memory in an International Consortium (COSMIC). (2020). Does parity matter in women’s risk of dementia? A COSMIC collaboration cohort study. BMC Medicine, 18(1), 210. https://doi.org/10.1186/s12916-020-01671-1

8. Baik, S. H., Kang, S., Son, S. M., & Mook-Jung, I. (2016). Microglia contributes to plaque growth by cell death due to uptake of amyloid β in the brain of Alzheimer’s disease mouse model. Glia, 64(12), 2274–2290. https://doi.org/10.1002/glia.23074

9. Barha, C. K., Lieblich, S. E., Chow, C., & Galea, L. A. M. (2015). Multiparity-induced enhancement of hippocampal neurogenesis and spatial memory depends on ovarian hormone status in middle age. Neurobiology of Aging, 36(8), 2391–2405. https://doi.org/10.1016/j.neurobiolaging.2015.04.007

10. Barnes, L. L., Wilson, R. S., Bienias, J. L., Schneider, J. A., Evans, D. A., & Bennett, D. A. (2005). Sex Differences in the Clinical Manifestations of Alzheimer Disease Pathology. Archives of General Psychiatry, 62(6), 685–691. https://doi.org/10.1001/archpsyc.62.6.685

11. Beeri, M. S., Rapp, M., Schmeidler, J., Reichenberg, A., Purohit, D. P., Perl, D. P., Grossman, H. T., Prohovnik, I., Haroutunian, V., & Silverman, J. M. (2009). Number of children is associated with neuropathology of Alzheimer’s disease in women. Neurobiology of Aging, 30(8), 1184–1191. https://doi.org/10.1016/j.neurobiolaging.2007.11.011

12. Belarbi, K., Arellano, C., Ferguson, R., Jopson, T., & Rosi, S. (2012). Chronic neuroinflammation impacts the recruitment of adult-born neurons into behaviorally relevant hippocampal networks. Brain, Behavior, and Immunity, 26(1), 18–23. https://doi.org/10.1016/j.bbi.2011.07.225

13. Berg, D. A., Bond, A. M., Ming, G., & Song, H. (2018). Radial glial cells in the adult dentate gyrus: What are they and where do they come from? F1000Research, 7, 277. https://doi.org/10.12688/f1000research.12684.1

14. Berger, A. (2000). Th1 and Th2 responses: What are they? BMJ: British Medical Journal, 321(7258), 424.

15. Boldrini, M., Fulmore, C. A., Tartt, A. N., Simeon, L. R., Pavlova, I., Poposka, V., Rosoklija, G. B., Stankov, A., Arango, V., Dwork, A. J., Hen, R., & Mann, J. J. (2018). Human Hippocampal Neurogenesis Persists throughout Aging. Cell Stem Cell, 22(4), 589-599.e5. https://doi.org/10.1016/j.stem.2018.03.015

16. Bolmont, T., Haiss, F., Eicke, D., Radde, R., Mathis, C. A., Klunk, W. E., Kohsaka, S., Jucker, M., & Calhoun, M. E. (2008). Dynamics of the Microglial/Amyloid Interaction Indicate a Role in Plaque Maintenance. The Journal of Neuroscience, 28(16), 4283. https://doi.org/10.1523/JNEUROSCI.4814-07.2008

17. Bozon, B., Kelly, A., Josselyn, S. A., Silva, A. J., Davis, S., & Laroche, S. (2003). MAPK, CREB and zif268 are all required for the consolidation of recognition memory. Philosophical Transactions of the Royal Society B: Biological Sciences, 358(1432), 805–814. https://doi.org/10.1098/rstb.2002.1224

18. Browne, T. C., McQuillan, K., McManus, R. M., O’Reilly, J.-A., Mills, K. H. G., & Lynch, M. A. (2013). IFN-γ Production by amyloid β-specific Th1 cells promotes microglial activation and increases plaque burden in a mouse model of Alzheimer’s disease. Journal of Immunology (Baltimore, Md.: 195), 190(5), 2241–2251. https://doi.org/10.4049/jimmunol.1200947

19. Castro-Portuguez, R., & Sutphin, G. L. (2020). Kynurenine pathway, NAD+ synthesis, and mitochondrial function: Targeting tryptophan metabolism to promote longevity and healthspan. Experimental Gerontology, 132, 110841. https://doi.org/10.1016/j.exger.2020.110841

20. Chatterjee, P., Goozee, K., Lim, C. K., James, I., Shen, K., Jacobs, K. R., Sohrabi, H. R., Shah, T., Asih, P. R., Dave, P., ManYan, C., Taddei, K., Lovejoy, D. B., Chung, R., Guillemin, G. J., & Martins, R. N. (2018). Alterations in serum kynurenine pathway metabolites in individuals with high neocortical amyloid-β load: A pilot study. Scientific Reports, 8. https://doi.org/10.1038/s41598-018-25968-7

21. Colombo, D., Serino, S., Tuena, C., Pedroli, E., Dakanalis, A., Cipresso, P., & Riva, G. (2017). Egocentric and allocentric spatial reference frames in aging: A systematic review. Neuroscience & Biobehavioral Reviews, 80, 605–621. https://doi.org/10.1016/j.neubiorev.2017.07.012

22. Colucci, M., Cammarata, S., Assini, A., Croce, R., Clerici, F., Novello, C., Mazzella, L., Dagnino, N., Mariani, C., & Tanganelli, P. (2006). The number of pregnancies is a risk factor for Alzheimer’s disease. European Journal of Neurology, 13(12), 1374–1377. https://doi.org/10.1111/j.1468-1331.2006.01520.x

23. Cui, J., Jothishankar, B., He, P., Staufenbiel, M., Shen, Y., & Li, R. (2014). Amyloid precursor protein mutation disrupts reproductive experience-enhanced normal cognitive development in a mouse model of Alzheimer’s disease. Molecular Neurobiology, 49(1), 103–112. https://doi.org/10.1007/s12035-013-8503-x

24. de Lange, A. G., Barth, C., Kaufmann, T., Anatürk, M., Suri, S., Ebmeier, K. P., & Westlye, L. T. (2020). The maternal brain: Region-specific patterns of brain aging are traceable decades after childbirth. Human Brain Mapping, 41(16), 4718–4729. https://doi.org/10.1002/hbm.25152

25. de Lange, A. G., Barth, C., Kaufmann, T., Maximov, I. I., van der Meer, D., Agartz, I., & Westlye, L. T. (2020). Women’s brain aging: Effects of sex-hormone exposure, pregnancies, and genetic risk for Alzheimer’s disease. Human Brain Mapping, 41(18), 5141–5150. https://doi.org/10.1002/hbm.25180

26. de Lange, A.-M. G., Kaufmann, T., van der Meer, D., Maglanoc, L. A., Alnæs, D., Moberget, T., Douaud, G., Andreassen, O. A., & Westlye, L. T. (2019). Population-based neuroimaging reveals traces of childbirth in the maternal brain. Proceedings of the National Academy of Sciences of the United States of America, 116(44), 22341–22346. https://doi.org/10.1073/pnas.1910666116

27. Duarte-Guterman, P., Albert, A. Y., Barha, C. K., & Galea, L. A. M. (2021). Sex influences the effects of APOE genotype and Alzheimer’s diagnosis on neuropathology and memory (p. 2020.06.25.20139980). https://doi.org/10.1101/2020.06.25.20139980

28. Duarte-Guterman, P., Richard, J. E., Lieblich, S. E., Eid, R. S., Lamers, Y., & Galea, L. a. M. (2023). Long-term cellular and molecular signatures of pregnancy in the adult and ageing brain (p. 2023.02.24.529879). bioRxiv. https://doi.org/10.1101/2023.02.24.529879

29. Dumurgier, J., Schraen, S., Gabelle, A., Vercruysse, O., Bombois, S., Laplanche, J.-L., Peoc’h, K., Sablonnière, B., Kastanenka, K. V., Delaby, C., Pasquier, F., Touchon, J., Hugon, J., Paquet, C., & Lehmann, S. (2015). Cerebrospinal fluid amyloid-β 42/40 ratio in clinical setting of memory centers: A multicentric study. Alzheimer’s Research & Therapy, 7(1), 30. https://doi.org/10.1186/s13195-015-0114-5

30. Eid, R. S., Chaiton, J. A., Lieblich, S. E., Bodnar, T. S., Weinberg, J., & Galea, L. A. M. (2019). Early and late effects of maternal experience on hippocampal neurogenesis, microglia, and the circulating cytokine milieu. Neurobiology of Aging, 78, 1–17. https://doi.org/10.1016/j.neurobiolaging.2019.01.021

31. Ekdahl, C. T., Claasen, J.-H., Bonde, S., Kokaia, Z., & Lindvall, O. (2003). Inflammation is detrimental for neurogenesis in adult brain. Proceedings of the National Academy of Sciences of the United States of America, 100(23), 13632–13637. https://doi.org/10.1073/pnas.2234031100

32. Ekonomou, A., Savva, G. M., Brayne, C., Forster, G., Francis, P. T., Johnson, M., Perry, E. K., Attems, J., Somani, A., Minger, S. L., & Ballard, C. G. (2015). Stage-Specific Changes in Neurogenic and Glial Markers in Alzheimer’s Disease. Biological Psychiatry, 77(8), 711–719. https://doi.org/10.1016/j.biopsych.2014.05.021

33. Eriksson, P. S., Perfilieva, E., Björk-Eriksson, T., Alborn, A.-M., Nordborg, C., Peterson, D. A., & Gage, F. H. (1998). Neurogenesis in the adult human hippocampus. Nature Medicine, 4(11), 1313–1317. https://doi.org/10.1038/3305

34. Ernst, A., Alkass, K., Bernard, S., Salehpour, M., Perl, S., Tisdale, J., Possnert, G., Druid, H., & Frisén, J. (2014). Neurogenesis in the striatum of the adult human brain. Cell, 156(5), 1072–1083. https://doi.org/10.1016/j.cell.2014.01.044

35. Fanselow, M. S., & Dong, H.-W. (2010). Are The Dorsal and Ventral Hippocampus functionally distinct structures? Neuron, 65(1), 7. https://doi.org/10.1016/j.neuron.2009.11.031

36. Farrer, L. A., Cupples, L. A., Haines, J. L., Hyman, B., Kukull, W. A., Mayeux, R., Myers, R. H., Pericak-Vance, M. A., Risch, N., & van Duijn, C. M. (1997). Effects of Age, Sex, and Ethnicity on the Association Between Apolipoprotein E Genotype and Alzheimer Disease: A Meta-analysis. JAMA, 278(16), 1349–1356. https://doi.org/10.1001/jama.1997.03550160069041

37. Ferretti, M. T., Iulita, M. F., Cavedo, E., Chiesa, P. A., Schumacher Dimech, A., Santuccione Chadha, A., Baracchi, F., Girouard, H., Misoch, S., Giacobini, E., Depypere, H., Hampel, H., & for the Women’s Brain Project and the Alzheimer Precision Medicine Initiative. (2018). Sex differences in Alzheimer disease—The gateway to precision medicine. Nature Reviews Neurology, 14(8), 457–469. https://doi.org/10.1038/s41582-018-0032-9

38. Floresco, S. B., Seamans, J. K., & Phillips, A. G. (1997). Selective Roles for Hippocampal, Prefrontal Cortical, and Ventral Striatal Circuits in Radial-Arm Maze Tasks With or Without a Delay. Journal of Neuroscience, 17(5), 1880–1890. https://doi.org/10.1523/JNEUROSCI.17-05-01880.1997

39. Franjic, D., Skarica, M., Ma, S., Arellano, J. I., Tebbenkamp, A. T. N., Choi, J., Xu, C., Li, Q., Morozov, Y. M., Andrijevic, D., Vrselja, Z., Spajic, A., Santpere, G., Li, M., Zhang, S., Liu, Y., Spurrier, J., Zhang, L., Gudelj, I.,…Sestan, N. (2022). Transcriptomic taxonomy and neurogenic trajectories of adult human, macaque, and pig hippocampal and entorhinal cells. Neuron, 110(3), 452–469.e14. https://doi.org/10.1016/j.neuron.2021.10.036

40. Gage, F. H. (2019). Adult neurogenesis in mammals. Science (New York, N.Y.), 364(6443), 827–828. https://doi.org/10.1126/science.aav6885

41. Galea, L. A. M., Roes, M. M., Dimech, C. J., Chow, C., Mahmoud, R., Lieblich, S. E., & Duarte-Guterman, P. (2018). Premarin has opposing effects on spatial learning, neural activation, and serum cytokine levels in middle-aged female rats depending on reproductive history. Neurobiology of Aging, 70, 291–307. https://doi.org/10.1016/j.neurobiolaging.2018.06.030

42. Håglin, S., Koch, E., Schäfer Hackenhaar, F., Nyberg, L., & Kauppi, K. (2023). APOE ɛ4, but not polygenic Alzheimer’s disease risk, is related to longitudinal decrease in hippocampal brain activity in non-demented individuals. Scientific Reports, 13(1), Article 1. https://doi.org/10.1038/s41598-023-35316-z

43. Haim, A., Julian, D., Albin-Brooks, C., Brothers, H. M., Lenz, K. M., & Leuner, B. (2017). A survey of neuroimmune changes in pregnant and postpartum female rats. Brain, Behavior, and Immunity, 59, 67–78. https://doi.org/10.1016/j.bbi.2016.09.026

44. Hohman, T. J., Dumitrescu, L., Barnes, L. L., Thambisetty, M., Beecham, G., Kunkle, B., Gifford, K. A., Bush, W. S., Chibnik, L. B., Mukherjee, S., Jager, P. L. D., Kukull, W., Crane, P. K., Resnick, S. M., Keene, C. D., Montine, T. J., Schellenberg, G. D., Haines, J. L., Zetterberg, H.,…Jefferson, A. L. (2018). Sex-Specific Association of Apolipoprotein E With Cerebrospinal Fluid Levels of Tau. JAMA Neurology, 75(8), 989–998. https://doi.org/10.1001/jamaneurol.2018.0821

45. Hollands, C., Tobin, M. K., Hsu, M., Musaraca, K., Yu, T.-S., Mishra, R., Kernie, S. G., & Lazarov, O. (2017). Depletion of adult neurogenesis exacerbates cognitive deficits in Alzheimer’s disease by compromising hippocampal inhibition. Molecular Neurodegeneration, 12(1), 64. https://doi.org/10.1186/s13024-017-0207-7

46. Irvine, K., Laws, K., Gale, T., & Kondel, K. (2012). Greater cognitive deterioration in women than men with Alzheimer’s disease: A meta analysis. Journal of Clinical and Experimental Neuropsychology, 34. https://doi.org/10.1080/13803395.2012.712676

47. Jang, H., Bae, J. B., Dardiotis, E., Scarmeas, N., Sachdev, P. S., Lipnicki, D. M., Han, J. W., Kim, T. H., Kwak, K. P., Kim, B. J., Kim, S. G., Kim, J. L., Moon, S. W., Park, J. H., Ryu, S.-H., Youn, J. C., Lee, D. Y., Lee, D. W., Lee, S. B.,…Kim, K. W. (2018). Differential effects of completed and incomplete pregnancies on the risk of Alzheimer disease. Neurology. https://doi.org/10.1212/WNL.0000000000006000

48. Jin, K., Galvan, V., Xie, L., Mao, X. O., Gorostiza, O. F., Bredesen, D. E., & Greenberg, D. A. (2004). Enhanced neurogenesis in Alzheimer’s disease transgenic (PDGF-APPSw,Ind) mice. Proceedings of the National Academy of Sciences of the United States of America, 101(36), 13363–13367. https://doi.org/10.1073/pnas.0403678101

49. Jin, K., Peel, A. L., Mao, X. O., Xie, L., Cottrell, B. A., Henshall, D. C., & Greenberg, D. A. (2004). Increased hippocampal neurogenesis in Alzheimer’s disease. Proceedings of the National Academy of Sciences of the United States of America, 101(1), 343–347. https://doi.org/10.1073/pnas.2634794100

50. Josephs, K. A., Dickson, D. W., Tosakulwong, N., Weigand, S. D., Murray, M. E., Petrucelli, L., Liesinger, A. M., Senjem, M. L., Spychalla, A. J., Knopman, D. S., Parisi, J. E., Petersen, R. C., Jack, C. R., & Whitwell, J. L. (2017). Rates of hippocampal atrophy and presence of post-mortem TDP-43 in patients with Alzheimer’s disease: A longitudinal retrospective study. The Lancet. Neurology, 16(11), 917–924. https://doi.org/10.1016/S1474-4422(17)30284-3

51. Kanai, M., Funakoshi, H., Takahashi, H., Hayakawa, T., Mizuno, S., Matsumoto, K., & Nakamura, T. (2009). Tryptophan 2,3-dioxygenase is a key modulator of physiological neurogenesis and anxiety-related behavior in mice. Molecular Brain, 2, 8. https://doi.org/10.1186/1756-6606-2-8

52. Kempermann, G., Gage, F. H., Aigner, L., Song, H., Curtis, M. A., Thuret, S., Kuhn, H. G., Jessberger, S., Frankland, P. W., Cameron, H. A., Gould, E., Hen, R., Abrous, D. N., Toni, N., Schinder, A. F., Zhao, X., Lucassen, P. J., & Frisén, J. (2018). Human Adult Neurogenesis: Evidence and Remaining Questions. Cell Stem Cell, 23(1), 25–30. https://doi.org/10.1016/j.stem.2018.04.004

53. Kinney, J. W., Bemiller, S. M., Murtishaw, A. S., Leisgang, A. M., Salazar, A. M., & Lamb, B. T. (2018). Inflammation as a central mechanism in Alzheimer’s disease. Alzheimer’s & Dementia: Translational Research & Clinical Interventions, 4, 575–590. https://doi.org/10.1016/j.trci.2018.06.014

54. Knoth, R., Singec, I., Ditter, M., Pantazis, G., Capetian, P., Meyer, R. P., Horvat, V., Volk, B., & Kempermann, G. (2010). Murine features of neurogenesis in the human hippocampus across the lifespan from 0 to 100 years. PloS One, 5(1), e8809. https://doi.org/10.1371/journal.pone.0008809

55. Korzhevskii, D. E., & Kirik, O. V. (2016). Brain Microglia and Microglial Markers. Neuroscience and Behavioral Physiology, 46(3), 284–290. https://doi.org/10.1007/s11055-016-0231-z

56. Koutseff, A., Mittelhaeuser, C., Essabri, K., Auwerx, J., & Meziane, H. (2014). Impact of the apolipoprotein E polymorphism, age and sex on neurogenesis in mice: Pathophysiological relevance for Alzheimer’s disease? Brain Research, 1542, 32–40. https://doi.org/10.1016/j.brainres.2013.10.003

57. Kwak, S. S., Washicosky, K. J., Brand, E., von Maydell, D., Aronson, J., Kim, S., Capen, D. E., Cetinbas, M., Sadreyev, R., Ning, S., Bylykbashi, E., Xia, W., Wagner, S. L., Choi, S. H., Tanzi, R. E., & Kim, D. Y. (2020). Amyloid-β42/40 ratio drives tau pathology in 3D human neural cell culture models of Alzheimer’s disease. Nature Communications, 11(1), Article 1. https://doi.org/10.1038/s41467-020-15120-3

58. Laws, K. R., Irvine, K., & Gale, T. M. (2016). Sex differences in cognitive impairment in Alzheimer’s disease. World Journal of Psychiatry, 6(1), 54–65. https://doi.org/10.5498/wjp.v6.i1.54

59. Lee, H., Price, J., Srivastava, D. P., & Thuret, S. (2023). In vitro characterization on the role of APOE polymorphism in human hippocampal neurogenesis. Hippocampus, 33(4), 322–346. https://doi.org/10.1002/hipo.23502

60. Lehmann, S., Delaby, C., Boursier, G., Catteau, C., Ginestet, N., Tiers, L., Maceski, A., Navucet, S., Paquet, C., Dumurgier, J., Vanmechelen, E., Vanderstichele, H., & Gabelle, A. (2018). Relevance of Aβ42/40 Ratio for Detection of Alzheimer Disease Pathology in Clinical Routine: The PLMR Scale. Frontiers in Aging Neuroscience, 10. https://www.frontiersin.org/articles/10.3389/fnagi.2018.00138

61. Leng, F., Hinz, R., Gentleman, S., Hampshire, A., Dani, M., Brooks, D. J., & Edison, P. (2023). Neuroinflammation is independently associated with brain network dysfunction in Alzheimer’s disease. Molecular Psychiatry, 28(3), Article 3. https://doi.org/10.1038/s41380-022-01878-z

62. Luckheeram, R. V., Zhou, R., Verma, A. D., & Xia, B. (2012). CD4+T Cells: Differentiation and Functions. Clinical and Developmental Immunology, 2012, 925135. https://doi.org/10.1155/2012/925135

63. Manning, E. N., Barnes, J., Cash, D. M., Bartlett, J. W., Leung, K. K., Ourselin, S., Fox, N. C., & Initiative, for the A. D. N. (2014). APOE ε4 Is Associated with Disproportionate Progressive Hippocampal Atrophy in AD. PLOS ONE, 9(5), e97608. https://doi.org/10.1371/journal.pone.0097608

64. Midttun, Ø., Hustad, S., Solheim, E., Schneede, J., & Ueland, P. M. (2005). Multianalyte Quantification of Vitamin B6 and B2 Species in the Nanomolar Range in Human Plasma by Liquid Chromatography-Tandem Mass Spectrometry. Clinical Chemistry, 51(7), 1206–1216.

65. Moreno-Jiménez, E. P., Flor-García, M., Terreros-Roncal, J., Rábano, A., Cafini, F., Pallas-Bazarra, N., Ávila, J., & Llorens-Martín, M. (2019). Adult hippocampal neurogenesis is abundant in neurologically healthy subjects and drops sharply in patients with Alzheimer’s disease. Nature Medicine, 25(4), 554–560. https://doi.org/10.1038/s41591-019-0375-9

66. Moser, M.-B., & Moser, E. I. (1998). Distributed Encoding and Retrieval of Spatial Memory in the Hippocampus. Journal of Neuroscience, 18(18), 7535–7542. https://doi.org/10.1523/JNEUROSCI.18-18-07535.1998

67. Mueller, S. G., Schuff, N., Yaffe, K., Madison, C., Miller, B., & Weiner, M. W. (2010). Hippocampal atrophy patterns in mild cognitive impairment and Alzheimer’s disease. Human Brain Mapping, 31(9), 1339–1347. https://doi.org/10.1002/hbm.20934

68. Neu, S. C., Pa, J., Kukull, W., Beekly, D., Kuzma, A., Gangadharan, P., Wang, L.-S., Romero, K., Arneric, S. P., Redolfi, A., Orlandi, D., Frisoni, G. B., Au, R., Devine, S., Auerbach, S., Espinosa, A., Boada, M., Ruiz, A., Johnson, S. C.,…Toga, A. W. (2017). Apolipoprotein E Genotype and Sex Risk Factors for Alzheimer’s Disease. JAMA Neurology, 74(10), 1178–1189. https://doi.org/10.1001/jamaneurol.2017.2188

69. Oxenkrug, G. F. (2010). Metabolic syndrome, age-associated neuroendocrine disorders, and dysregulation of tryptophan-kynurenine metabolism. Annals of the New York Academy of Sciences, 1199, 1–14. https://doi.org/10.1111/j.1749-6632.2009.05356.x

70. Parizkova, M., Lerch, O., Moffat, S. D., Andel, R., Mazancova, A. F., Nedelska, Z., Vyhnalek, M., Hort, J., & Laczó, J. (2018). The effect of Alzheimer’s disease on spatial navigation strategies. Neurobiology of Aging, 64, 107–115. https://doi.org/10.1016/j.neurobiolaging.2017.12.019

71. Posillico, C. K., & Schwarz, J. M. (2016). An investigation into the effects of antenatal stressors on the postpartum neuroimmune profile and depressive-like behaviors. Behavioural Brain Research, 298(0 0), 218–228. https://doi.org/10.1016/j.bbr.2015.11.011

72. Puri, T. A., Richard, J. E., & Galea, L. A. M. (2023). Beyond sex differences: Short-and long-term effects of pregnancy on the brain. Trends in Neurosciences, 46(6), 459–471. https://doi.org/10.1016/j.tins.2023.03.010

73. Riedel, B. C., Thompson, P. M., & Brinton, R. D. (2016). Age, APOE and sex: Triad of risk of Alzheimer’s disease. The Journal of Steroid Biochemistry and Molecular Biology, 160, 134–147. https://doi.org/10.1016/j.jsbmb.2016.03.012

74. Sanai, N., Nguyen, T., Ihrie, R. A., Mirzadeh, Z., Tsai, H.-H., Wong, M., Gupta, N., Berger, M. S., Huang, E., Garcia-Verdugo, J.-M., Rowitch, D. H., & Alvarez-Buylla, A. (2011). Corridors of migrating neurons in the human brain and their decline during infancy. Nature, 478(7369), 382–386. https://doi.org/10.1038/nature10487

75. Sohn, D., Shpanskaya, K., Lucas, J. E., Petrella, J. R., Saykin, A. J., Tanzi, R. E., Samatova, N. F., & Doraiswamy, P. M. (2018). Sex Differences in Cognitive Decline in Subjects with High Likelihood of Mild Cognitive Impairment due to Alzheimer’s disease. Scientific Reports, 8(1), 7490. https://doi.org/10.1038/s41598-018-25377-w

76. Sorgdrager, F. J. H., Vermeiren, Y., Van Faassen, M., van der Ley, C., Nollen, E. A. A., Kema, I. P., & De Deyn, P. P. (2019). Age- and disease-specific changes of the kynurenine pathway in Parkinson’s and Alzheimer’s disease. Journal of Neurochemistry, 151(5), 656–668. https://doi.org/10.1111/jnc.14843

77. Sorrells, S. F., Paredes, M. F., Cebrian-Silla, A., Sandoval, K., Qi, D., Kelley, K. W., James, D., Mayer, S., Chang, J., Auguste, K. I., Chang, E. F., Gutierrez, A. J., Kriegstein, A. R., Mathern, G. W., Oldham, M. C., Huang, E. J., Garcia-Verdugo, J. M., Yang, Z., & Alvarez-Buylla, A. (2018). Human hippocampal neurogenesis drops sharply in children to undetectable levels in adults. Nature, 555(7696), Article 7696. https://doi.org/10.1038/nature25975

78. Spalding, K. L., Bergmann, O., Alkass, K., Bernard, S., Salehpour, M., Huttner, H. B., Boström, E., Westerlund, I., Vial, C., Buchholz, B. A., Possnert, G., Mash, D. C., Druid, H., & Frisén, J. (2013). Dynamics of hippocampal neurogenesis in adult humans. Cell, 153(6), 1219–1227. https://doi.org/10.1016/j.cell.2013.05.002

79. Stalder, M., Phinney, A., Probst, A., Sommer, B., Staufenbiel, M., & Jucker, M. (1999). Association of Microglia with Amyloid Plaques in Brains of APP23 Transgenic Mice. The American Journal of Pathology, 154(6), 1673–1684.

80. Steiner, B., Klempin, F., Wang, L., Kott, M., Kettenmann, H., & Kempermann, G. (2006). Type-2 cells as link between glial and neuronal lineage in adult hippocampal neurogenesis. Glia, 54(8), 805– 814. https://doi.org/10.1002/glia.20407

81. Streit, W. J. (2005). Microglia and neuroprotection: Implications for Alzheimer’s disease. Brain Research. Brain Research Reviews, 48(2), 234–239. https://doi.org/10.1016/j.brainresrev.2004.12.013

82. Tai, L. M., Bilousova, T., Jungbauer, L., Roeske, S. K., Youmans, K. L., Yu, C., Poon, W. W., Cornwell, L. B., Miller, C. A., Vinters, H. V., Van Eldik, L. J., Fardo, D. W., Estus, S., Bu, G., Gylys, K. H., & Ladu, M. J. (2013). Levels of soluble apolipoprotein E/amyloid-β (Aβ) complex are reduced and oligomeric Aβ increased with APOE4 and Alzheimer disease in a transgenic mouse model and human samples. The Journal of Biological Chemistry, 288(8), 5914–5926. https://doi.org/10.1074/jbc.M112.442103

83. Taxier, L. R., Philippi, S. M., Fleischer, A. W., York, J. M., LaDu, M. J., & Frick, K. M. (2022). APOE4 homozygote females are resistant to the beneficial effects of 17β-estradiol on memory and CA1 dendritic spine density in the EFAD mouse model of Alzheimer’s disease. Neurobiology of Aging, 118, 13–24. https://doi.org/10.1016/j.neurobiolaging.2022.06.005

84. Tensaouti, Y., Stephanz, E. P., Yu, T.-S., & Kernie, S. G. (2018). ApoE Regulates the Development of Adult Newborn Hippocampal Neurons. Eneuro, 5(4), ENEURO.0155-18.2018. https://doi.org/10.1523/ENEURO.0155-18.2018

85. Thal, D. R., Holzer, M., Rüb, U., Waldmann, G., Günzel, S., Zedlick, D., & Schober, R. (2000). Alzheimer-Related τ-Pathology in the Perforant Path Target Zone and in the Hippocampal Stratum Oriens and Radiatum Correlates with Onset and Degree of Dementia. Experimental Neurology, 163(1), 98–110. https://doi.org/10.1006/exnr.2000.7380

86. Tobin, M. K., Musaraca, K., Disouky, A., Shetti, A., Bheri, A., Honer, W. G., Kim, N., Dawe, R. J., Bennett, D. A., Arfanakis, K., & Lazarov, O. (2019). Human Hippocampal Neurogenesis Persists in Aged Adults and Alzheimer’s Disease Patients. Cell Stem Cell, 24(6), 974–982.e3. https://doi.org/10.1016/j.stem.2019.05.003

87. Town, T., Vendrame, M., Patel, A., Poetter, D., DelleDonne, A., Mori, T., Smeed, R., Crawford, F., Klein, T., Tan, J., & Mullan, M. (2002). Reduced Th1 and enhanced Th2 immunity after immunization with Alzheimer’s β-amyloid1–42. Journal of Neuroimmunology, 132(1), 49–59. https://doi.org/10.1016/S0165-5728(02)00307-7

88. Wiener, J. M., de Condappa, O., Harris, M. A., & Wolbers, T. (2013). Maladaptive bias for extrahippocampal navigation strategies in aging humans. The Journal of Neuroscience: The Official Journal of the Society for Neuroscience, 33(14), 6012–6017. https://doi.org/10.1523/JNEUROSCI.0717-12.2013

89. Wolk, D. A., Dickerson, B. C., & Initiative, the A. D. N. (2010). Apolipoprotein E (APOE) genotype has dissociable effects on memory and attentional–executive network function in Alzheimer’s disease. Proceedings of the National Academy of Sciences, 107(22), 10256–10261. https://doi.org/10.1073/pnas.1001412107

90. Wu, M. D., Hein, A. M., Moravan, M. J., Shaftel, S. S., Olschowka, J. A., & O’Banion, M. K. (2012). Adult murine hippocampal neurogenesis is inhibited by sustained IL-1β and not rescued by voluntary running. Brain, Behavior, and Immunity, 26(2), 292–300. https://doi.org/10.1016/j.bbi.2011.09.012

91. Wu, W., Nicolazzo, J. A., Wen, L., Chung, R., Stankovic, R., Bao, S. S., Lim, C. K., Brew, B. J., Cullen, K. M., & Guillemin, G. J. (2013). Expression of tryptophan 2,3-dioxygenase and production of kynurenine pathway metabolites in triple transgenic mice and human Alzheimer’s disease brain. PloS One, 8(4), e59749. https://doi.org/10.1371/journal.pone.0059749

92. Yagi, S., Chow, C., Lieblich, S. E., & Galea, L. A. M. (2016). Sex and strategy use matters for pattern separation, adult neurogenesis, and immediate early gene expression in the hippocampus. Hippocampus, 26(1), 87–101. https://doi.org/10.1002/hipo.22493

93. Yagi, S., Splinter, J. E. J., Tai, D., Wong, S., Wen, Y., & Galea, L. A. M. (2020). Sex Differences in Maturation and Attrition of Adult Neurogenesis in the Hippocampus. ENeuro, 7(4), ENEURO.0468-19.2020. https://doi.org/10.1523/ENEURO.0468-19.2020

94. Zhou, Y., Su, Y., Li, S., Kennedy, B. C., Zhang, D. Y., Bond, A. M., Sun, Y., Jacob, F., Lu, L., Hu, P., Viaene, A. N., Helbig, I., Kessler, S. K., Lucas, T., Salinas, R. D., Gu, X., Chen, H. I., Wu, H., Kleinman, J. E.,…Song, H. (2022). Molecular landscapes of human hippocampal immature neurons across lifespan. Nature, 1–7. https://doi.org/10.1038/s41586-022-04912-w

95. Zunszain, P. A., Anacker, C., Cattaneo, A., Choudhury, S., Musaelyan, K., Myint, A. M., Thuret, S., Price, J., & Pariante, C. M. (2012). Interleukin-1β: A New Regulator of the Kynurenine Pathway Affecting Human Hippocampal Neurogenesis. Neuropsychopharmacology, 37(4), Article 4. https://doi.org/10.1038/npp.2011.277

96. Zwilling, D., Huang, S.-Y., Sathyasaikumar, K. V., Notarangelo, F. M., Guidetti, P., Wu, H.-Q., Lee, J., Truong, J., Andrews-Zwilling, Y., Hsieh, E. W., Louie, J. Y., Wu, T., Scearce-Levie, K., Patrick, C., Adame, A., Giorgini, F., Moussaoui, S., Laue, G., Rassoulpour, A.,…Muchowski, P. J. (2011). Kynurenine 3-monooxygenase inhibition in blood ameliorates neurodegeneration. Cell, 145(6), 863–874. https://doi.org/10.1016/j.cell.2011.05.020

